# Unravelling Immune Cell Signatures: CIBERSORTx-assisted Construction of Signature Matrix from Single-Cell Data

**DOI:** 10.1101/2024.05.05.592045

**Authors:** Madhulika Verma

## Abstract

The immune system’s intricate orchestration is pivotal in combating infections and diseases, often leaving discernible signatures within circulating blood cells. Peripheral blood mononuclear cells (PBMCs), comprising diverse immune cell populations, serve as crucial indicators of immune system status and its responses to various conditions, including cancer. While traditional bulk metrics pose challenges in dissecting specific immune cell functionalities, advancements in single-cell technologies offer unprecedented insights into the dynamic activities of immune cell populations.

In this study, we analyzed single-cell data from droplet sequencing to delineate immune cell types and subtypes within PBMCs. Employing the CIBERSORTx tool, we constructed a signature matrix to comprehensively represent significant cell populations within tissue. Through iterative optimization and minimization of the condition number based on marker genes, we aimed to enhance the robustness and stability of the gene signature matrices, enabling scalable investigation of novel or poorly understood phenotypic states in bulk tissue gene expression profiles. The resultant matrix consisted of 14 immune cell types represented by 275 gene having significant highest expression for their respective cell types and the least in other cells.

This approach facilitates precise characterization of immune cell populations and their responses to diverse diseases, contributing to a deeper understanding of immunological processes and paving the way for targeted therapeutic interventions.

## 1. Introduction

The immune system plays a pivotal role in defending the body against infections and diseases. While accessing tissue from the site of illness can be challenging, analyzing blood samples offers a viable alternative to assess immune system status and its involvement in various conditions (Lawlor et al., 2021; Sen et al., 2017; Schmiedel et al., 2018). Immune cells circulating in the bloodstream reflect the body’s health status through distinct signatures, which can undergo changes in cellular structure, function, gene expression, or DNA (Kotliarov et al., 2020; Banchereau et al., 2016).

Peripheral blood mononuclear cells (PBMCs) are integral to the immune system’s ability to combat diseases. However, due to the diverse nature of PBMCs, comprising various immune cell types, investigating specific cell populations’ functionality often relies on bulk metrics (Khan & Kaihara, 2019). Examples of PBMCs include monocytes, dendritic cells (DCs), and lymphocytes, characterized by their round nuclei (Bardou et al., 2014). PBMCs, being a key component of the immune system, play critical roles in immunological responses to diverse diseases, notably cancer (Bloodworth et al., 2016; Boniface et al., 2009; Cader et al., 2016).

Advancements in single-cell technologies have enabled precise characterization of immune cell populations in clinical samples containing mixed cell populations. These technologies offer insights into the dynamic cellular activities of immune populations, such as cytotoxic T lymphocytes, dendritic cells, and B lymphocytes, facilitating better understanding of immunological diseases (Papalexi & Satija, 2017; Pereira et al., 2020; Shalek et al., 2014; Horns et al., 2020).

In this study, we utilized PBMC single-cell data obtained through droplet sequencing to comprehensively identify and classify various immune cell types and subtypes. Subsequently, distinct clusters for each cell type, characterized by significant marker genes, were delineated. These clusters were then leveraged to construct a signature matrix employing the CIBERSORTx tool (Steen et al., 2020).

CIBERSORTx is a tool that was primarily used for the deconvolution of bulk RNA-seq data, in other words, for quantifying the abundance of different cell types in a bulk RNA-seq data, using a “signature matrix” which consisted of cell-specific signatures provided by the user or the one created by authors using the microarray data, named as “LM22”. Now it also has a utility that enables the construction of a “signature matrix” without the need for laborious sorting trials, thereby encompassing the entire significant populations of cells within tissue, using single-cell RNA sequencing (scRNA-seq). This approach allows for the investigation of novel or poorly understood phenotypic states in bulk tissue gene expression profiles on a massive scale. The signature matrix captures cell-specific expression signatures, which are crucial for understanding immune cell dynamics.

The condition number, which measures the sensitivity of the function’s output value to changes in input arguments, was utilized to assess the robustness of the signature matrix. By iteratively adjusting the number of marker genes included in the matrix and evaluating the condition number, we aimed to minimize its value to enhance the matrix’s stability. Specifically, the 2-norm condition number, calculated using the kappa function in R (Newman et al., 2015), was employed to quantify the stability of the signature matrix.

## 2. Materials and Methods

### 2.1. Data

The data was collected from the 10x genomics website (https://www.10xgenomics.com/datasets/frozen-pbm-cs-donor-c-1-standard-1-1-0) and sequenced using 10x genomics chromium technology (Home Page - 10x Genomics, n.d.. In this technique, the Gel bead in Emulsion (GEM) method is used to put single cells in the shape of droplets (which is why this is one of the droplet-based sequencing techniques), labelling each gel bead with oligonucleotides unique to each bead and having a length of 10 bp, also known as barcodes or UMIs (unique molecular identifiers). Additionally, they have sequencing primers and adapters and an immobile attached oligo-dT of 30 bp.

The datasets were presented as a raw counts sparse grid array. (pre-analyzed, primary analysis done using Cell Ranger 1.1.0), with 9519 cells in the column and 32738 genes as rows.

### 2.2. Analysis

**Figure 1.**
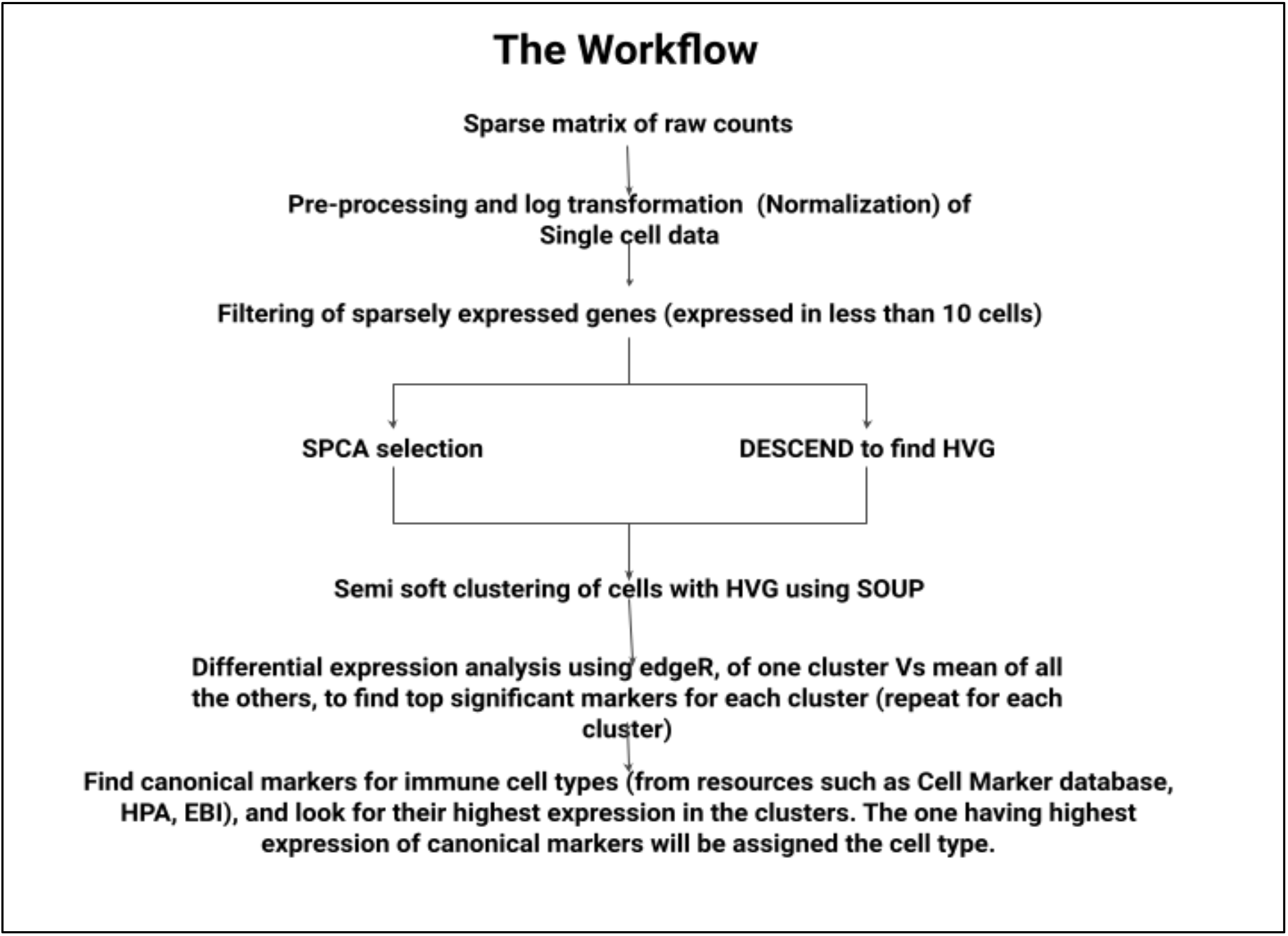
The pipeline for the analysis of the scRNA seq derived PBMC data.

### 2.3. Pre-processing and HVG selection

The data was first “pre-processed,” which means it was TPM-normalized and log-transformed. Then, the highly variable gene selection method was used to find the most informative genes. For the clustering step, only those informative genes which showed maximum variation across cells in the data were chosen. So, the gene selection step not only picked the genes that changed a lot, but it also eliminated genes that were present in very few cells (because they were caught by chance or noise). Elimination of the housekeeping genes and the genes which were expressed uniformly throughout was done at this stage as they won’t be useful in the following step of cluster separation.

“Semi-soft clustering of single-cell data (SOUP)” (Zhu et al., 2018) was used for gene selection and clustering steps. This tool provides two methods for selecting highly variable and informative genes. First is the sparse PCA algorithm (SPCA), which is a lasso (elastic net)-based PCA method specialized for sparse loadings such as single-cell datasets (Witten et al., 2009). PCA is used for dimensionality reduction of the data by selecting a smaller number of variables that retain most of the sample’s information. It often happens that in the case of classic PCA, the input variables are typically combined linearly to form the principal components. This makes the result unpredictable and makes it hard to determine what the principal components mean since they are all not zero. This issue is resolved with sparse PCA by looking for linear combinations that use only a few input variables. Thus, in the case of gene expression data, each variable corresponds to a specific gene. With only a small number of genes, sparse PCA can generate a principal component so that the analysis can be narrowed down to just these genes. This method is better suited for single-cell datasets because it finds linear combinations while efficiently utilizing a small number of input variables, unlike the classic PCA method, which is typically a linear combination of all input variables. Also, the sparse PCA can retain consistency, which the classic PCA cannot.

The second method of gene selection to reduce noise and filter out less informative genes is the DESCEND algorithm (Wang et al., 2018). DESCEND (“deconvolution of single-cell expression distribution”, Wang et al., 2018) utilizes Gini coefficients to provide accurate estimations of measures based on distributions, such as the actual gene expression variance and the likelihood that true gene expression is positive. While preserving the genes that are more plausible to serve as cell subtype identifiers, low-variance genes are filtered away to reduce noise. The coefficient can take on values between 0 and 1, with 0 denoting the least possible variation and 1 denoting highly variable expression throughout. Thus, the greater the Gini index, the more distributed the data.

Together, these two methods choose a group of informative genes that shows high variance in their expressions in different cells and are also informative. These genes are then further subjected to clustering.

### 2.4. Clustering

The next step is clustering the cells using the selected genes. SOUP is a method for semi-soft clustering scRNA datasets that fills a void in characterizing cells with ambiguous labels, such as those that transition from one type of cell to another (Zhu et al., 2018). The SOUP procedure begins with identifying a collection of pure cells, followed by estimating the non-negative membership matrix. Considering that the data has (i) Pure cells belonging to a common group that requires a hard cluster allocation and (ii) Developmental cells (transitional cells) that are shifting among different cell types and should be assigned lenient allocations, SOUP first finds K different clusters (or cell subsets) from data and then assigns membership scores to mixed cells for each cluster K, forming a “membership matrix”. The membership matrix’s every row has positive numbers that add up to one. These numbers show how many cells are in each of the K clusters. Specifically, a type k non-transitional cell has a membership score of 1 and zeros elsewhere. Here, hard clustering is utilized for this data. An elbow plot (optimal K) was used to figure out K, and Scater (McCarthy et al., 2017) was used to make PCA, TSNE (t-distributed stochastic neighbour embeddings), and UMAP (Uniform manifold approximation and projection) plots. These plots are typically used to visualize single-cell data in low-dimensional space. They are also known as “non-linear dimensional reduction methods.”

### 2.5. Differential Expression Analysis of the Clusters

After the clustering, edgeR was used for in-depth analysis of the clusters In edgeR analysis, each cluster (including all of the cell types in the cluster) was treated as a sample, similar to other conventional differential expression analyses. The Differential analysis was carried out to determine potential key markers for each cluster. The contrasts were made to compare one cluster and the mean of all the remaining clusters. This procedure yielded a list of the most significant genes for each cluster. Then, various databases like the Human Protein Atlas, the CellMarker Database, and the EBI were used to find the standard markers of immune cells. The top genes from each cluster were searched for in the above-mentioned databases. According to the search, the cell subsets were assigned to the cluster based on which immune cell their top genes were markers for.

### 2.6. Construction of the Signature Matrix

After obtaining the matrix of cell types and their marker genes, the tool CIBERSORTx was implemented to construct a custom signature matrix. A .txt or .tsv file containing a tab-delimited matrix of scRNA-seq expression data, with rows representing genes and columns representing single-cell transcriptomic data, makes up the input file. It was checked to make sure the sum of all the genes expressed in a given cell wasn’t zero. The first column of the file was filled with gene names. Redundant genes were removed. Also, each cell in row 1 was given a cell phenotype, such as “CD8 T cell” or “B cell,” and it was made sure that there were at least three cells of each phenotype. The highly expressed G marker genes from every cell subset were integrated into a signature matrix BG, and CIBERSORT sorted significant genes for each cell type subset in decreasing order of fold change in comparison to other cell subsets. The algorithm preserved the signature matrix, identifying the one with the minimum condition number after iterating G from 50 to 200 across all subgroups. To avoid genes expressed on non-hematopoietic cell types confusing the findings of deconvolution, the option to filter out non-hematopoietic genes were activated. After we obtained the signature matrix for the immune cells, visualizations were drawn by means of a heatmap and hierarchical clustering to ensure the correct groupings and sub-groupings.

## 3. Results and Discussion

### 3.1. Pre-processing

The downloaded PBMC data had 9519 cells in the column and 32738 genes as rows. Since the data for these genes were raw counts, it was TPM normalized and log transformed. To find out the existing grouping in this data, single-cell linear and non-linear dimensionality reduction approaches were used. We utilized PCA, TSNE, and UMAP plots to view the data in lower dimensions (figure 2.).

**Figure 2.**
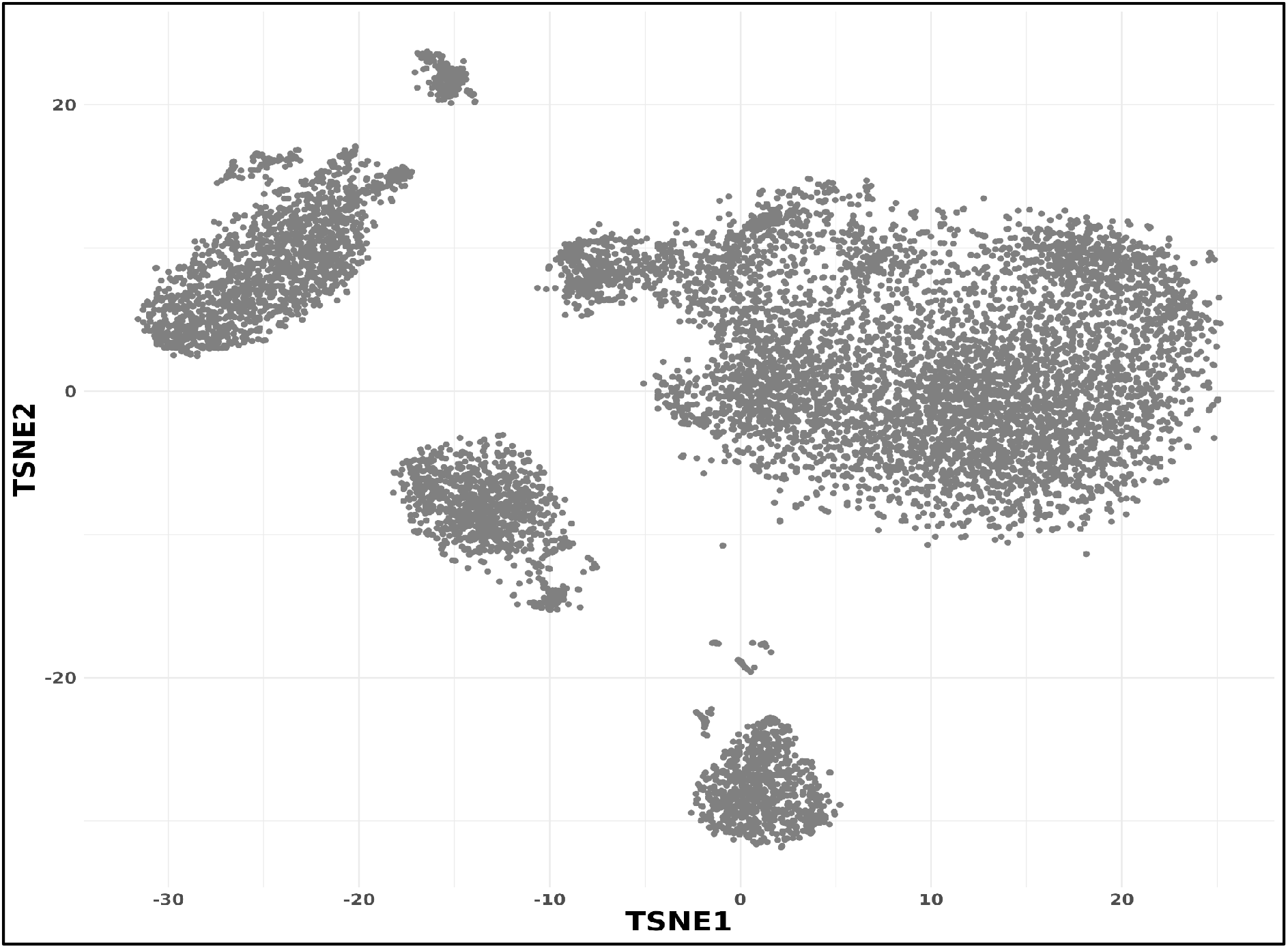
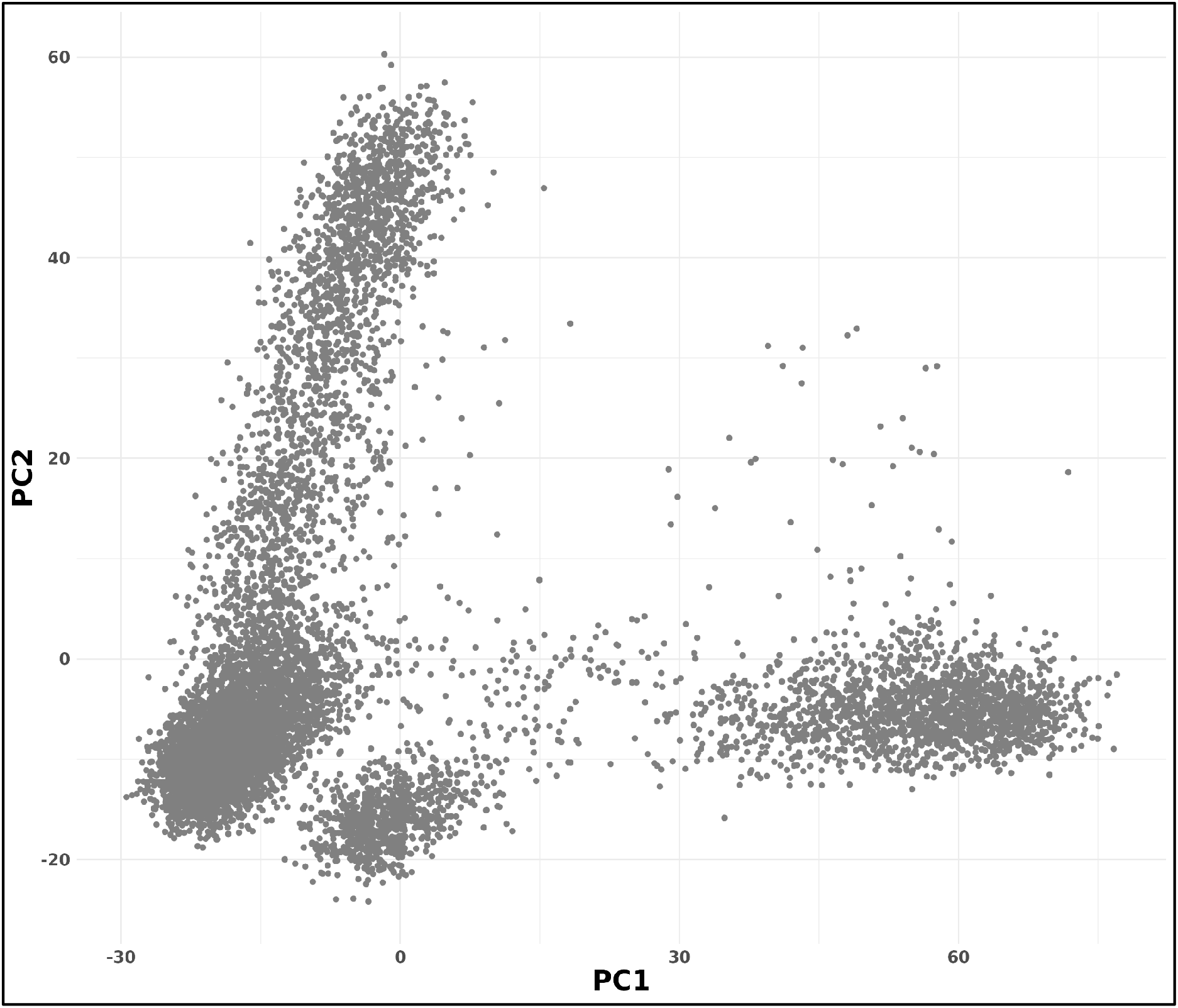
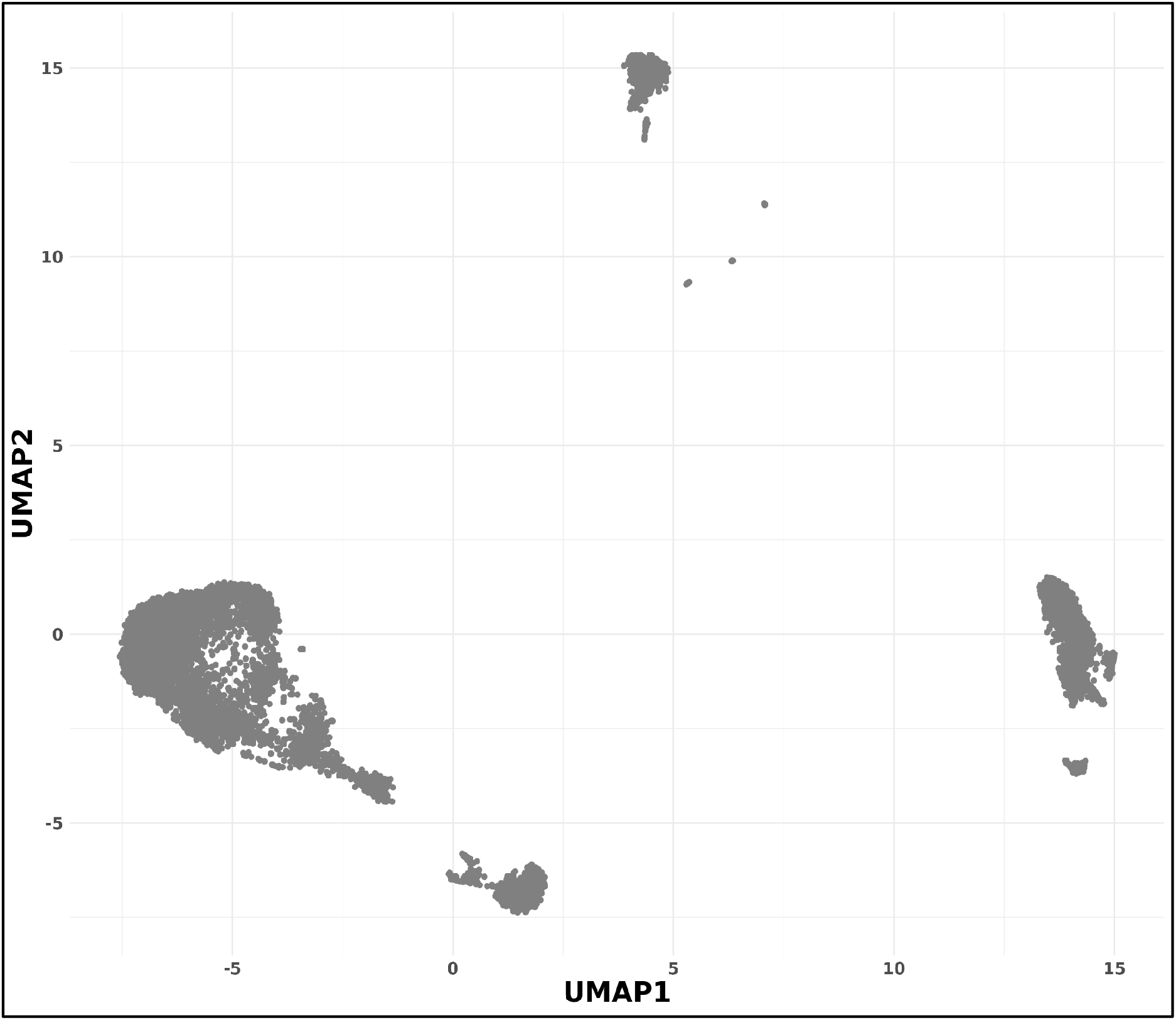
TSNE, PCA, and UMAP of PBMC data. These plots show that the data is divided into at least eight distinct groups.

### 3.2. HVG selection

Single-cell data usually requires clustering of the samples to find out the structure present in them, but before clustering, it is important to filter out the non-informative genes. In the first stage, we removed the sparsely expressed genes or genes whose expression was detected in less than ten cells. Then DESCEND, and SPCA was applied to find out highly variable genes from the data. The DESCEND selected 609 genes, and the SPCA selected 158 genes using default parameters.

Fig. 3. Shows the distribution and variance of the genes. The curve was made using quantile smooth regression (the default quantile is 0.5), demonstrating the correlation between the mean of deconvolved distributions and the Gini coefficient of the genes. The Gini distance from the curve is used to figure out each gene’s dispersion score. This score is normalized by the gene’s SE (squared error). Genes with a Gini coefficient and a normalized score higher than a threshold (the default is 10) are marked as having a lot of variation as shown in red. Total of 678 genes out of 32738 were identified as highly variable and the analysis was further continued with this set of highly variable genes only (along with all the cells). SOUP selected 678 genes when the gene selection was performed using default parameters. However, the number of selected genes may be increased or decreased when the parameters such as “threshold” and “sumabs” are changed in the gene selection step. The “threshold” is the cutoff for the Gini index of DESCEND. Fewer genes are picked when the threshold is higher. Similarly, “sumabs” is a measurement of the sparsity of genes in SPCA, between 1/sqrt(number of genes) and 1. Smaller values result in sparser results, hence fewer selected genes.

**Figure 3.**
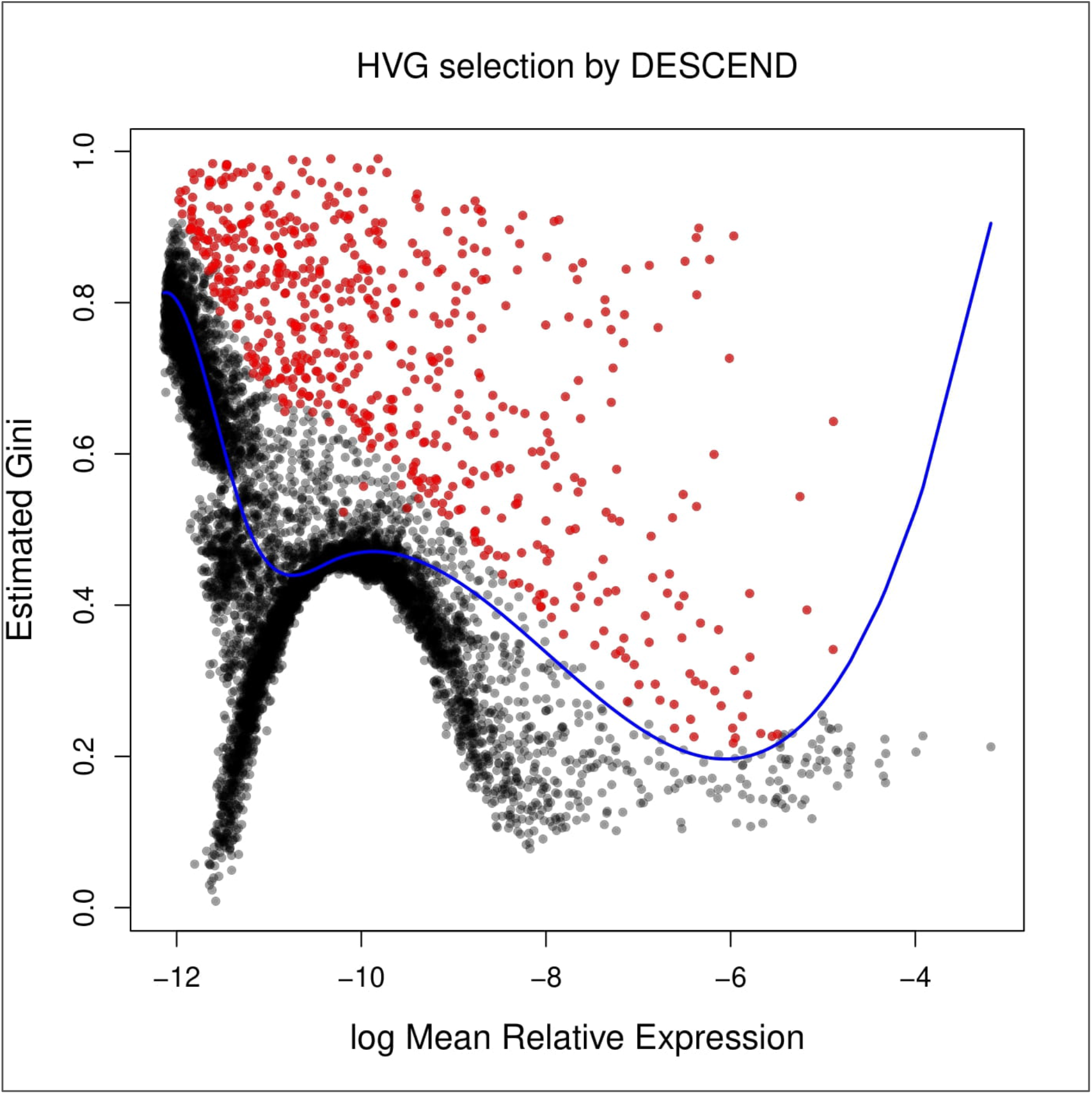
HVG selected by DESCEND. The distribution and variance of the genes. The curve was made using quantile smooth regression (the default quantile is 0.5), showing the relationship between the mean of deconvolved distributions and Gini across genes. The Gi distance from the curve is used to figure out each gene’s dispersion score. This score is normalized by the gene’s squared error). Genes with a Gini coefficient and a normalized score higher than a threshold (the default is 10) are marked as having a lot of variation. The data points in red show this. Out of 32738 genes, 678 were designated as highly variable.

### 3.3. Clustering

Clustering techniques were performed as a next step after getting highly variable genes. SOUP allows users to specify the value of K for clustering. For the identification of optimal K in our data, we considered the elbow plot. To choose the ideal K, the number of clusters, SOUP offers a cross-validation process. Specifically, using 10-fold cross-validation, users can search across K > 2, > 10, (or more), and the mistakes are averaged over nCV repeats (Zhu et al., 2018). Cross-validation errors were visualized in the log scale when comparing several K values and discovered that K = 8 had the lowest cross-validation error. This suggests that there are at least eight groups in our PBMC data. The selected highly variable genes are able to segregate the data into 8 distinct groups. If the number of genes selected as highly variable is increased or decreased, then, maybe the number of groupings in our data also changes because when the data was visualized in PCA, t-SNE, and UMAP more than 8 groupings of cells were found. These other groupings may be subtypes of cells for example activated cells or resting cells and so on.

**Figure 4.**
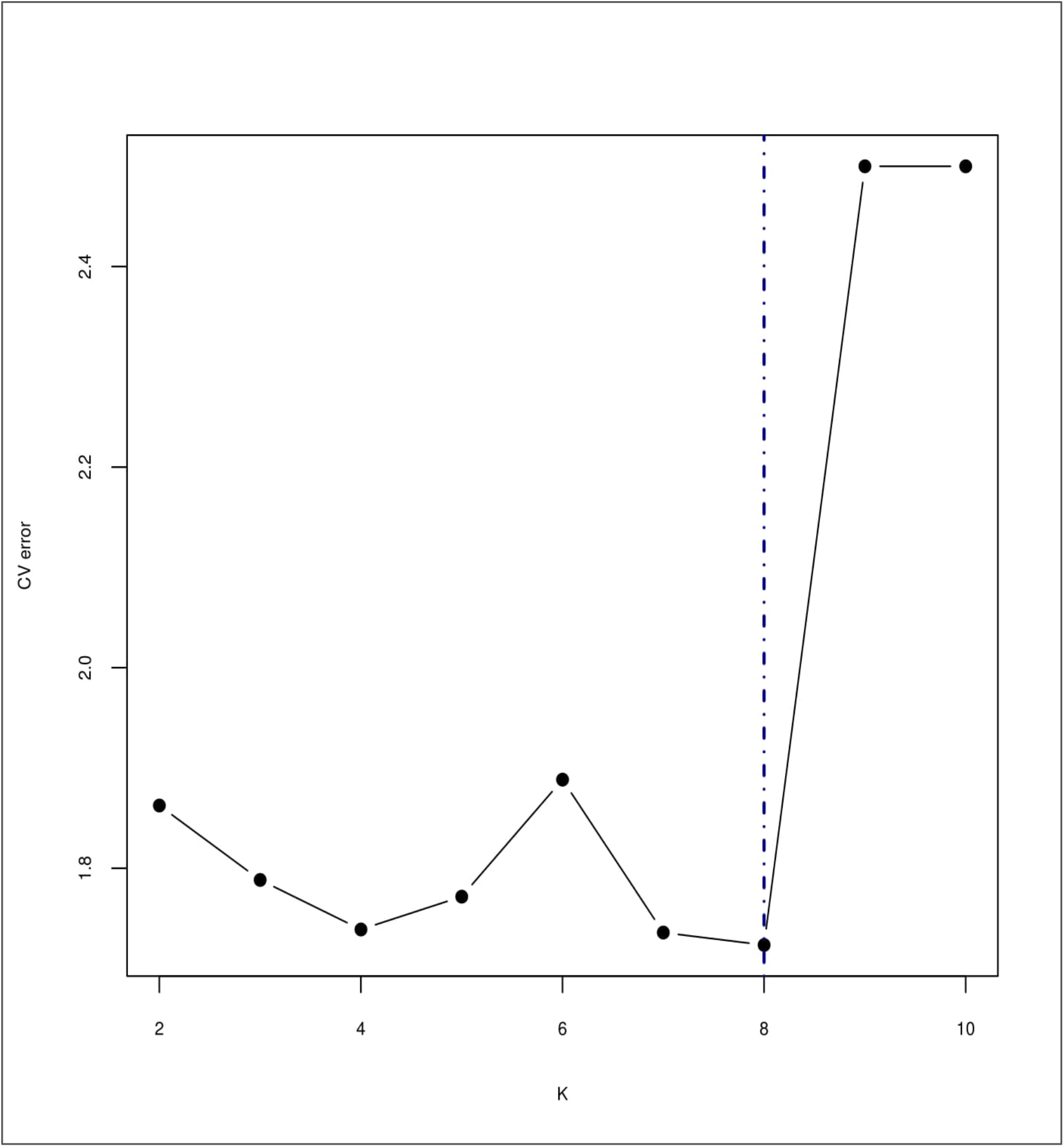
Elbow plot showing the optimal K in the PBMC data. The elbow falls the lowest cross-validation error to 8, which shows that our data has at least 8 clusters.

Therefore, we continued to cluster the data by increasing K until we obtained clusters with fewer than five cells. Thus, we tested K = 10 and increased K until this condition was met, stopping at K = 15.

**Table 1.** Table showing clustering results of K = 15.

| Cl. | 1 | 2 | 3 | 4 | 5 | 6 | 7 | 8 | 9 | 10 | 11 | 13 | 14 | 15 |
| --- | --- | --- | --- | --- | --- | --- | --- | --- | --- | --- | --- | --- | --- | --- |
| No. off cells | 4793 | 8 | 494 | 980 | 256 | 436 | 66 | 481 | 622 | 211 | 40 | 20 | 364 | 748 |

We further proceeded with K = 15 results, and the clusters were annotated on the above visualizations.

**Figure 5.**
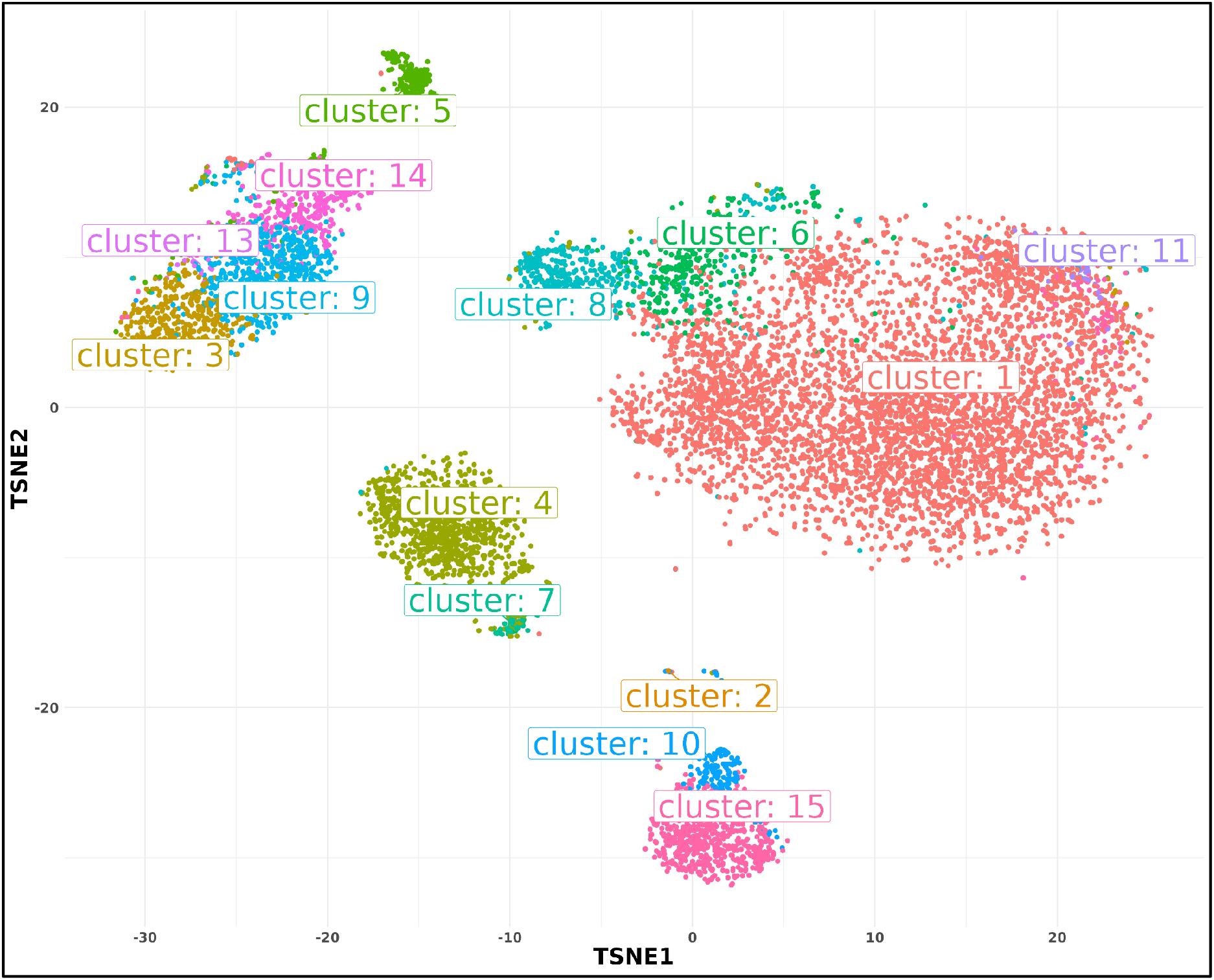
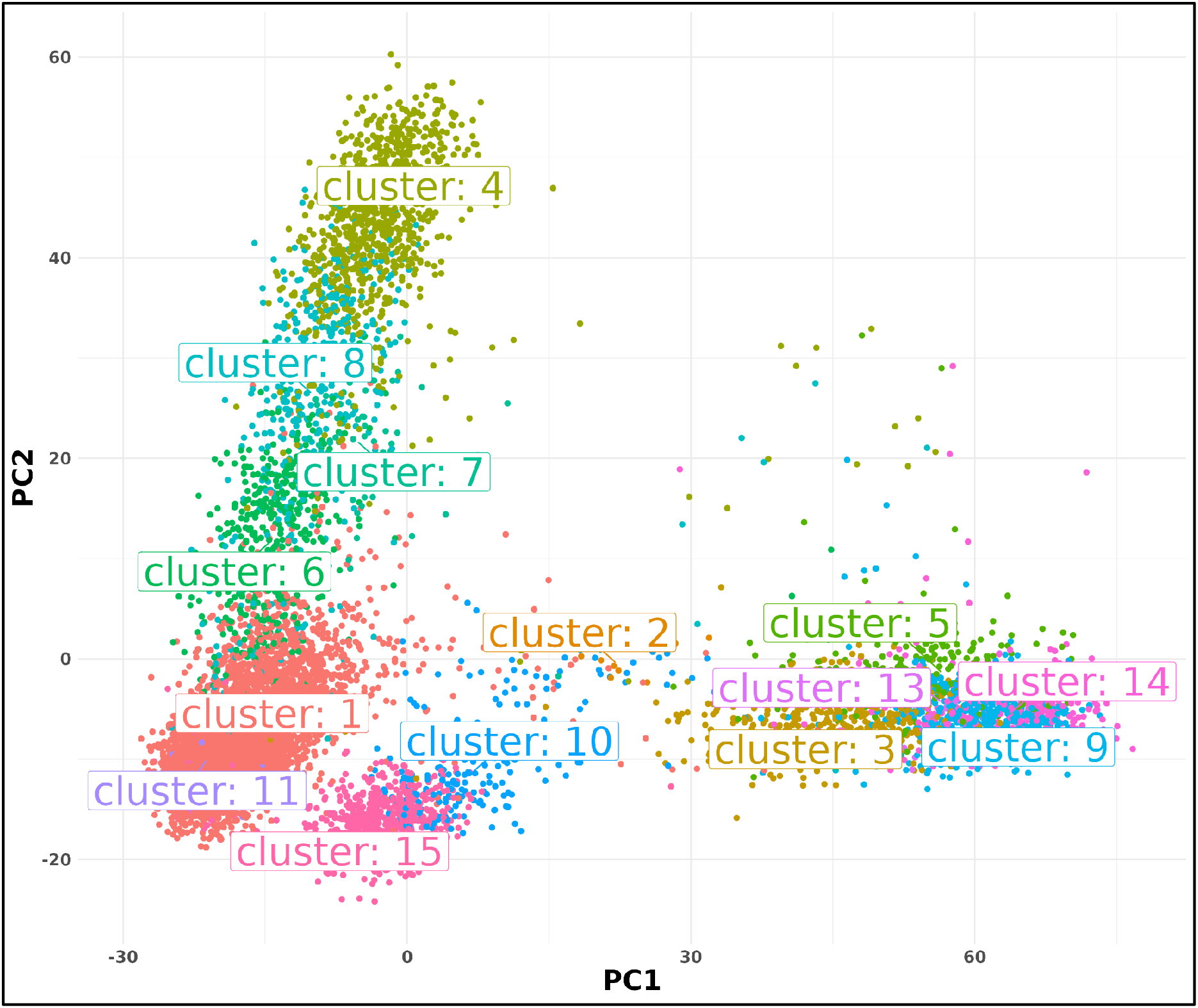
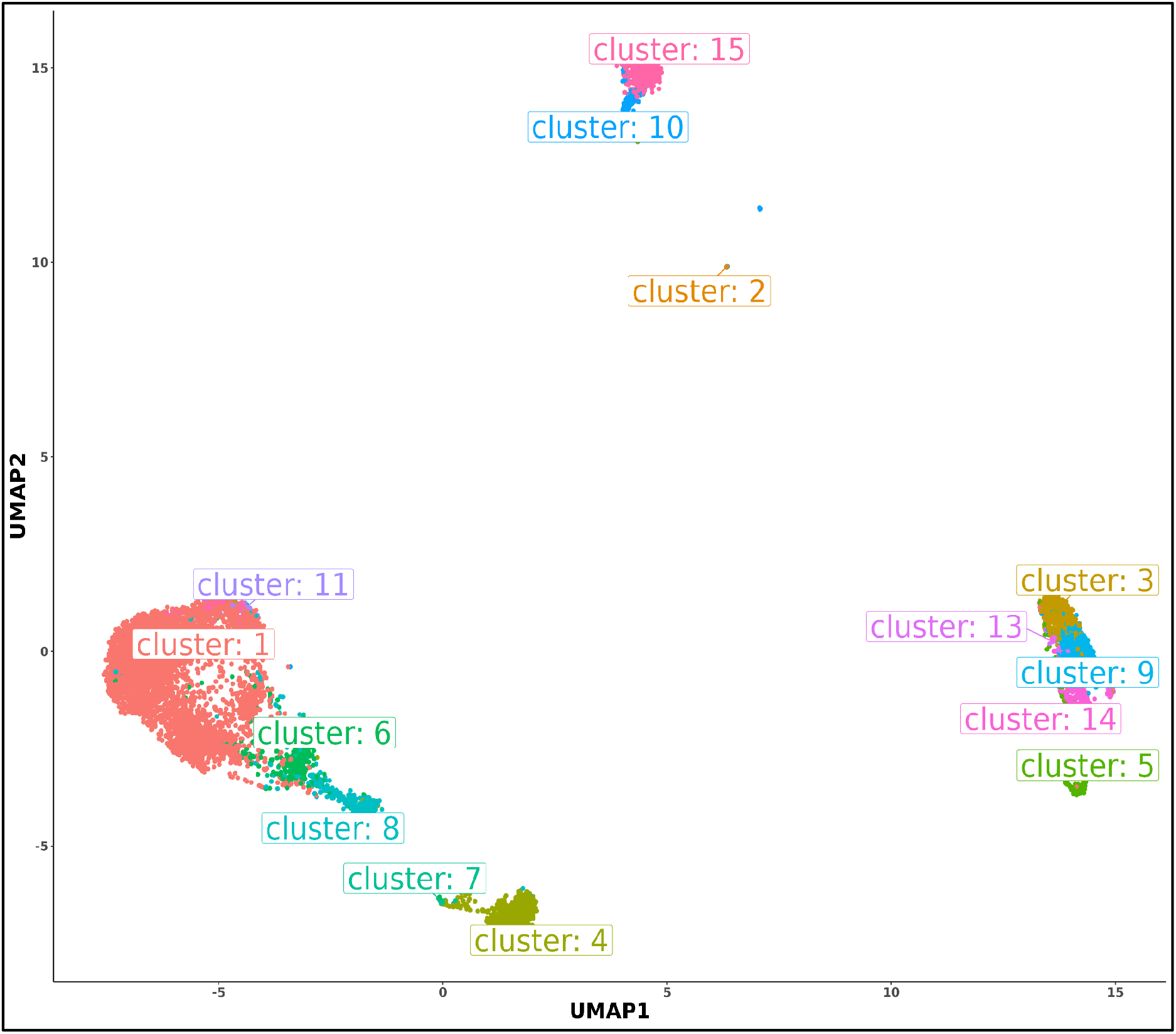
TSNE, PCA, and UMAP plot showing clusters identified in the PBMC data plots showing clusters identified in PBMC data.

### 3.4. Differential expression analysis of the clusters

After the clustering methods r, differential expression analysis was done on the clustered groups. The goal was to find the top significant genes that could separate a cluster from the rest of the clusters. For this purpose, we considered using the classic differential expression analysis tool, edgeR (Robinson et al., 2010). EdgeR favors the raw integer read counts and finds a scaling (normalization) factor for every sample to normalize the read counts for different library sizes (sample-specific effect) (Robinson et al., 2010). The TMM (Trimmed Mean of M-values) (Robinson et al., 2010) between-sample normalization approach was used for the normalization. TMM presupposes that most of the genes do not express themselves differently. The scaling factors were then used to calculate the actual library sizes. CR or Cox-Reid profile-adjusted likelihood (CR) is used in the EdgeR approach to estimate dispersion for trials with many components. After getting estimates of dispersion and fitting negative binomial generalized linear models, we moved on to the testing steps to find differential expressions using the likelihood ratio test.

A design matrix is necessary for the GLM technique in order to specify the treatment conditions. To create the design matrix, we used the “**model**.**matrix”** function. From the groupings of clusters, we made a model matrix in which each cell had a 1 under the cluster where it was found and a 0 everywhere else, as shown below:

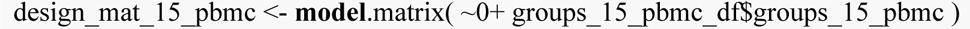

Using the QL F-test, we looked for genes that showed significant differential expression. The “makeContrasts” function was used to specify the contrast of interest. The comparisons were set up so that one cluster was put up against the average of every other cluster, as shown below:

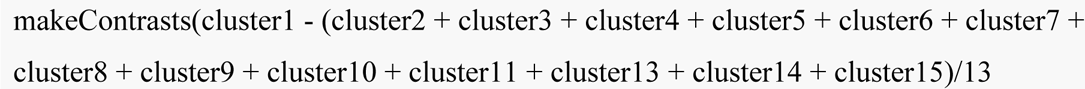

This testing process provided the significant genes for each cluster (positively and negatively expressed) (Table 2. top 10 significant genes from each cluster).

**Figure 6.**
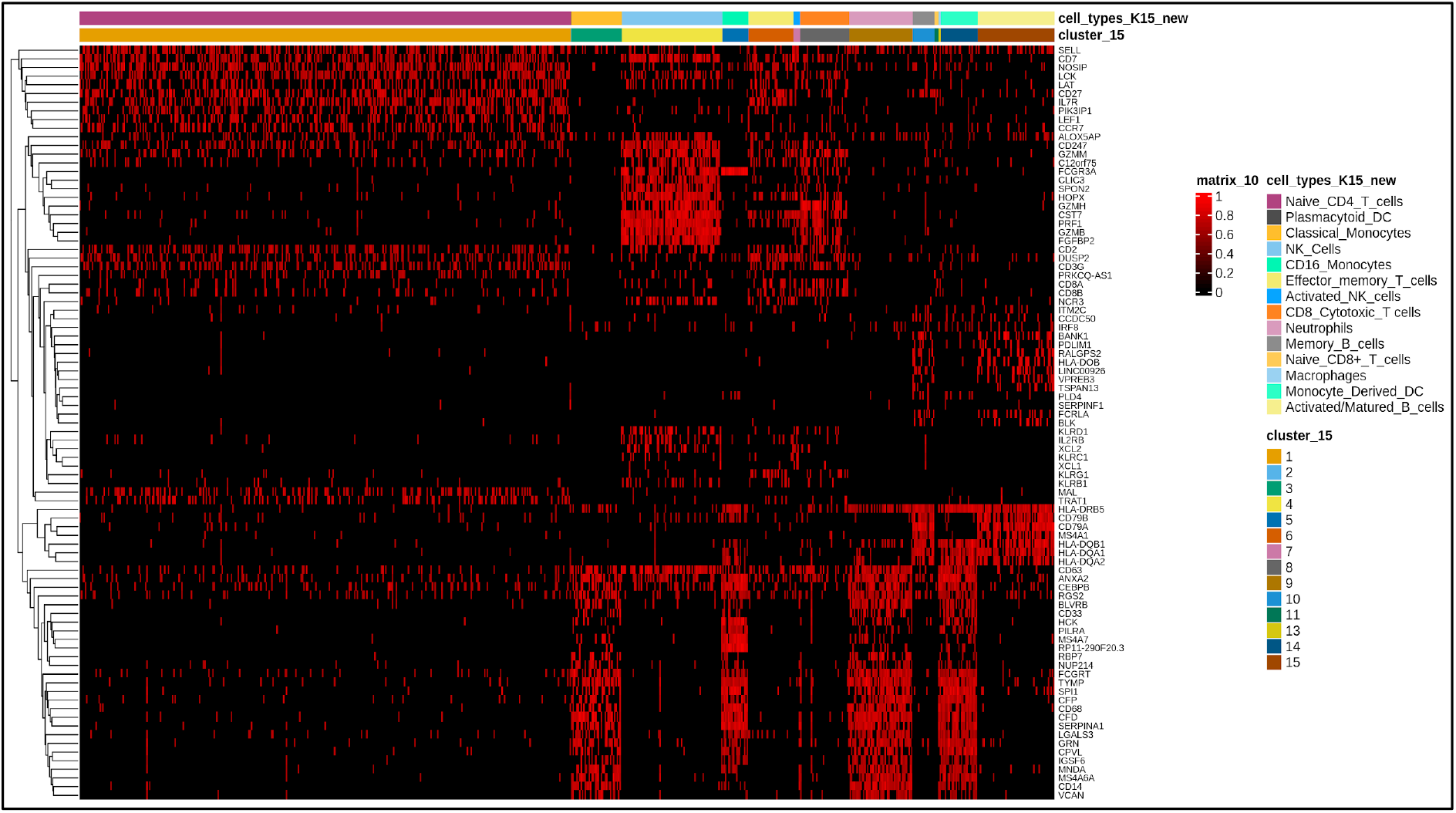
Heatmap of top 10 significant genes from each cluster and their differential expression across the clusters.

**Table 2.**
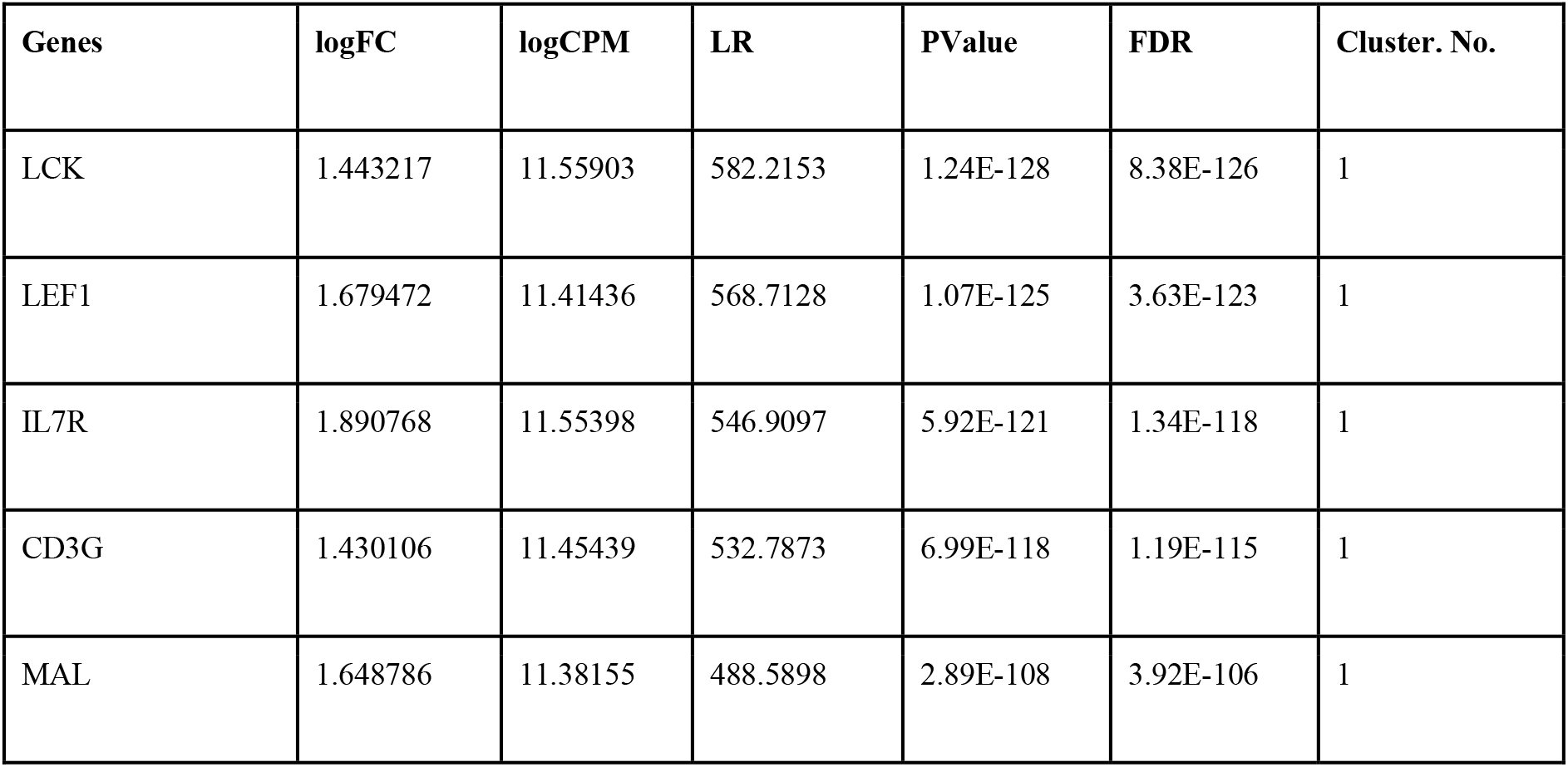
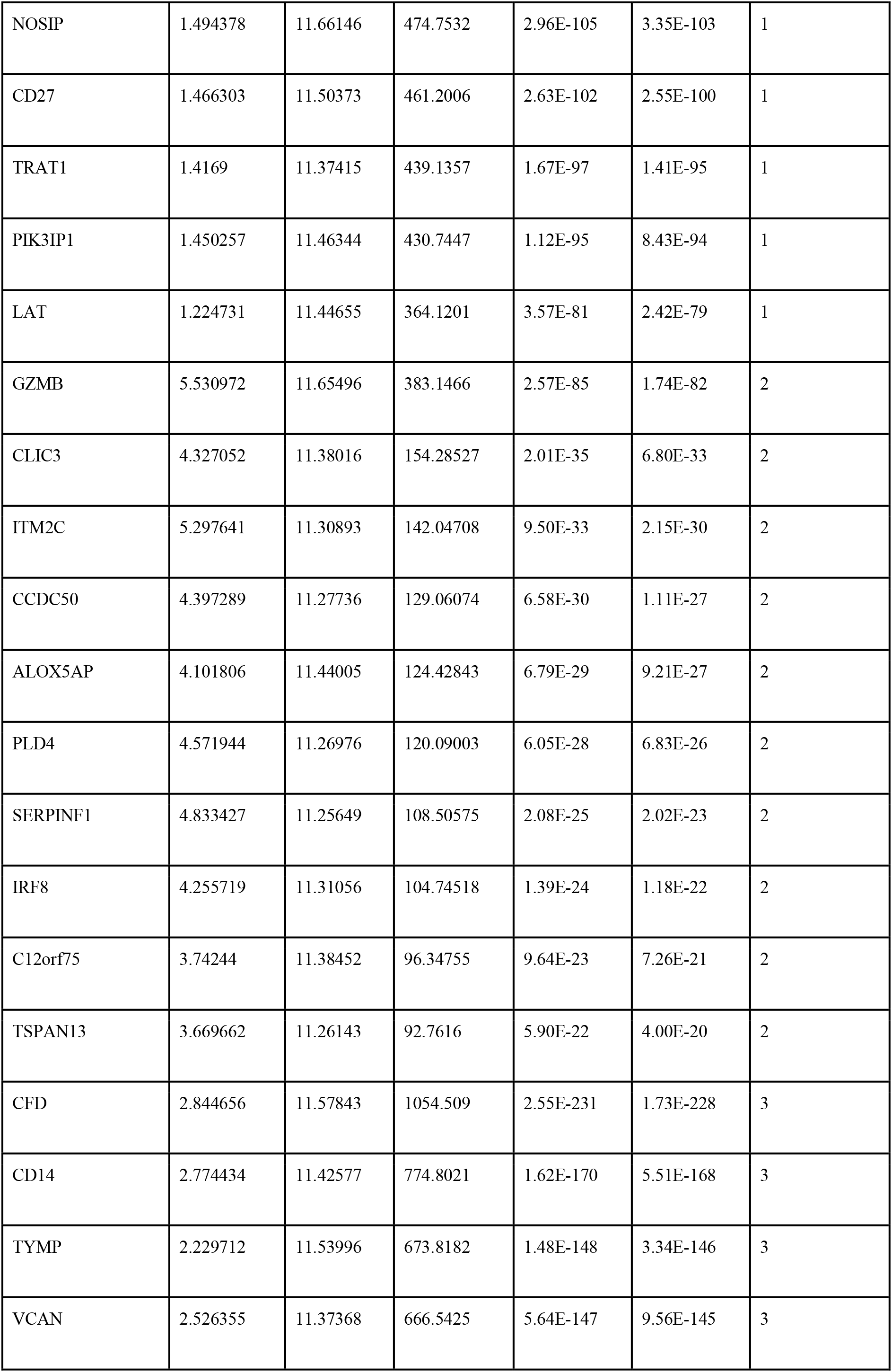
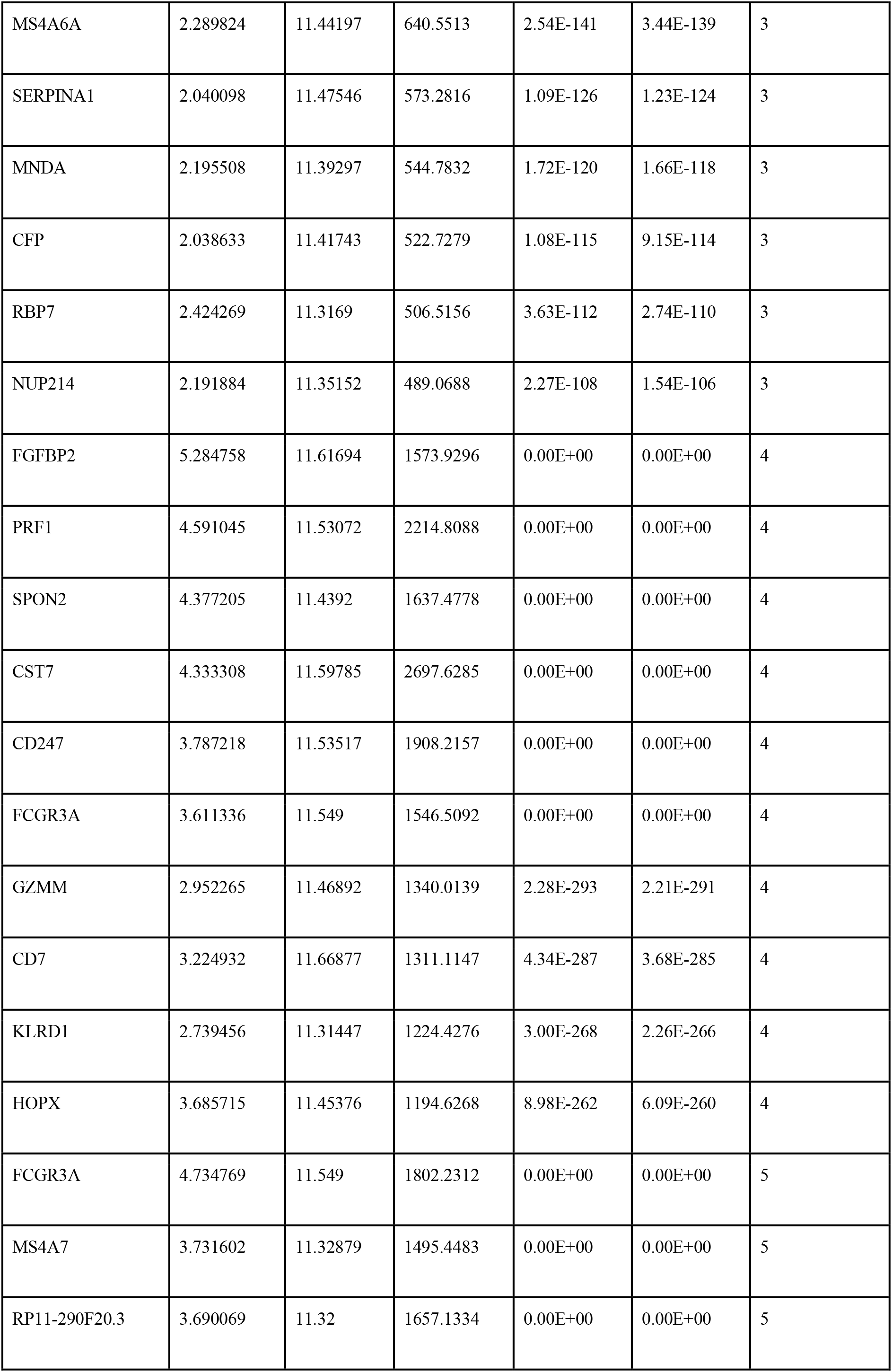
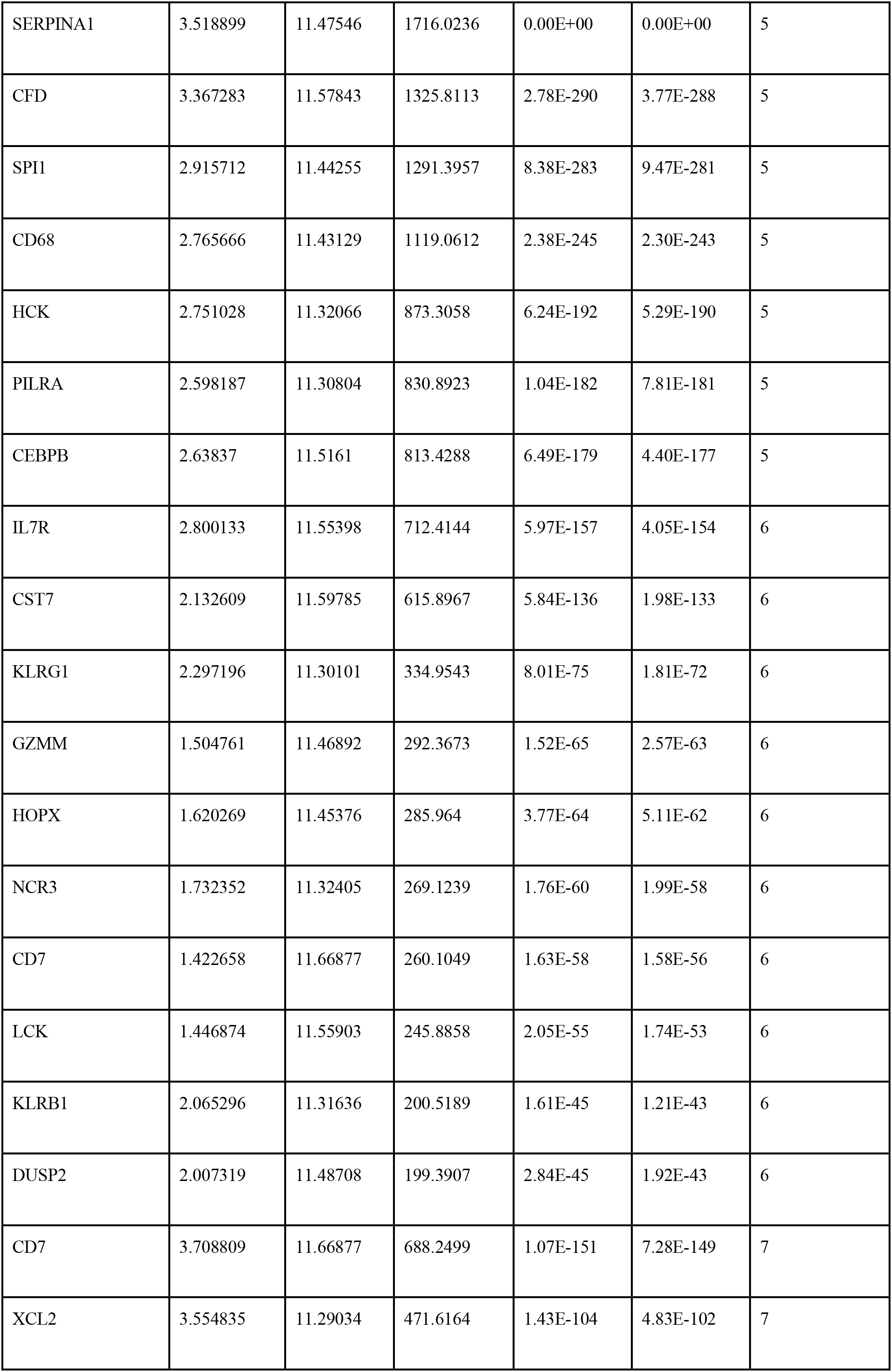
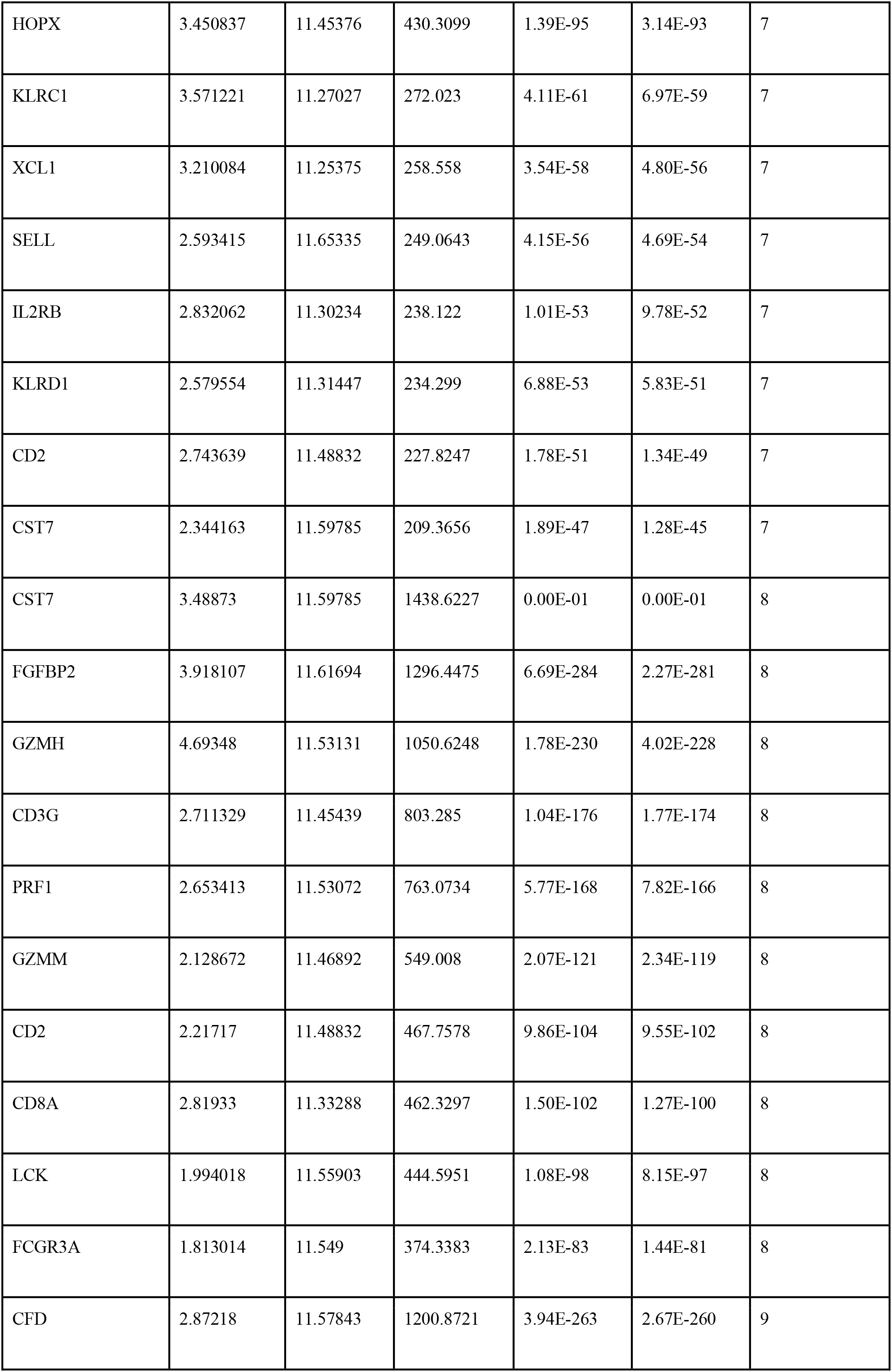
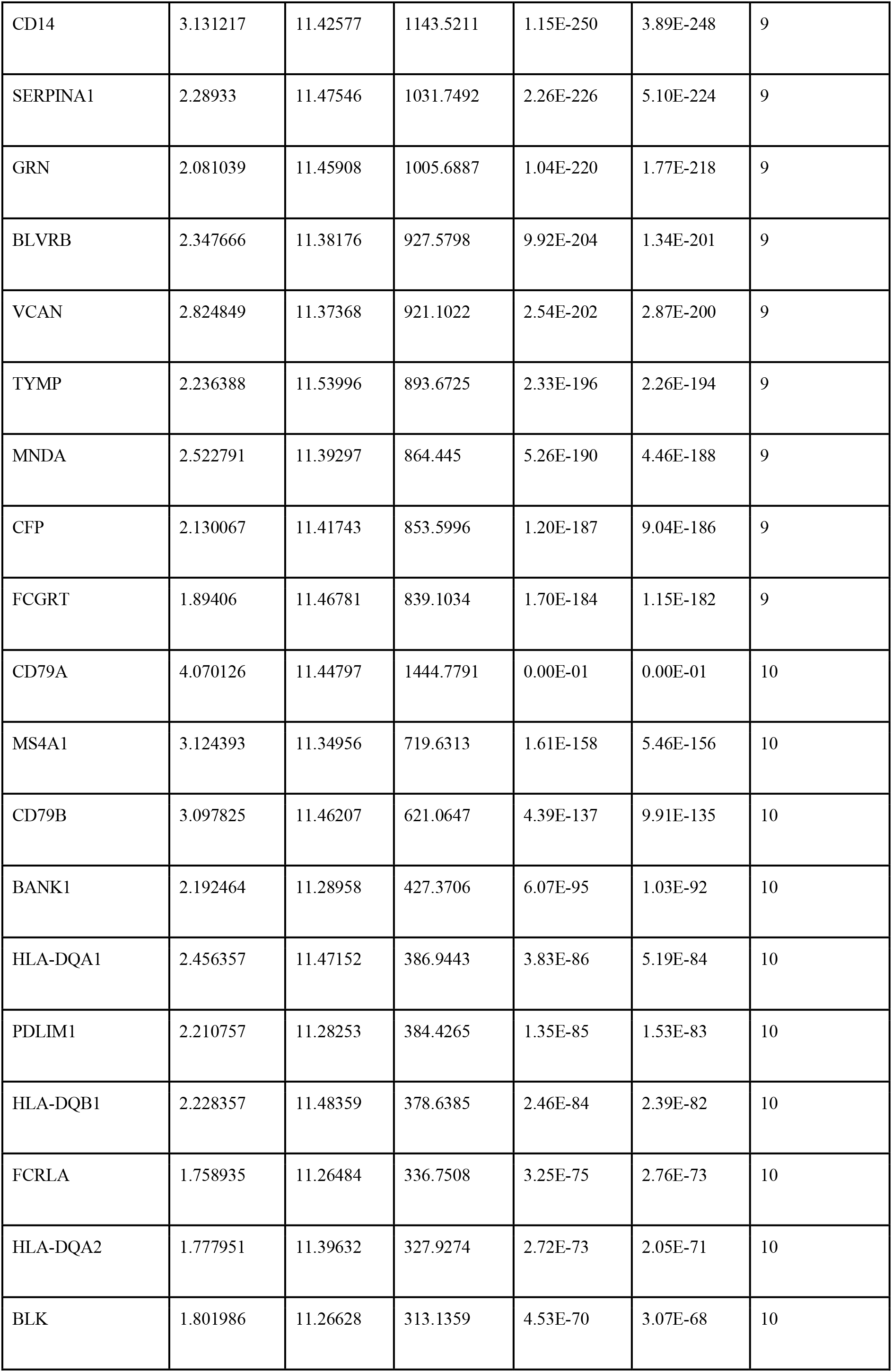
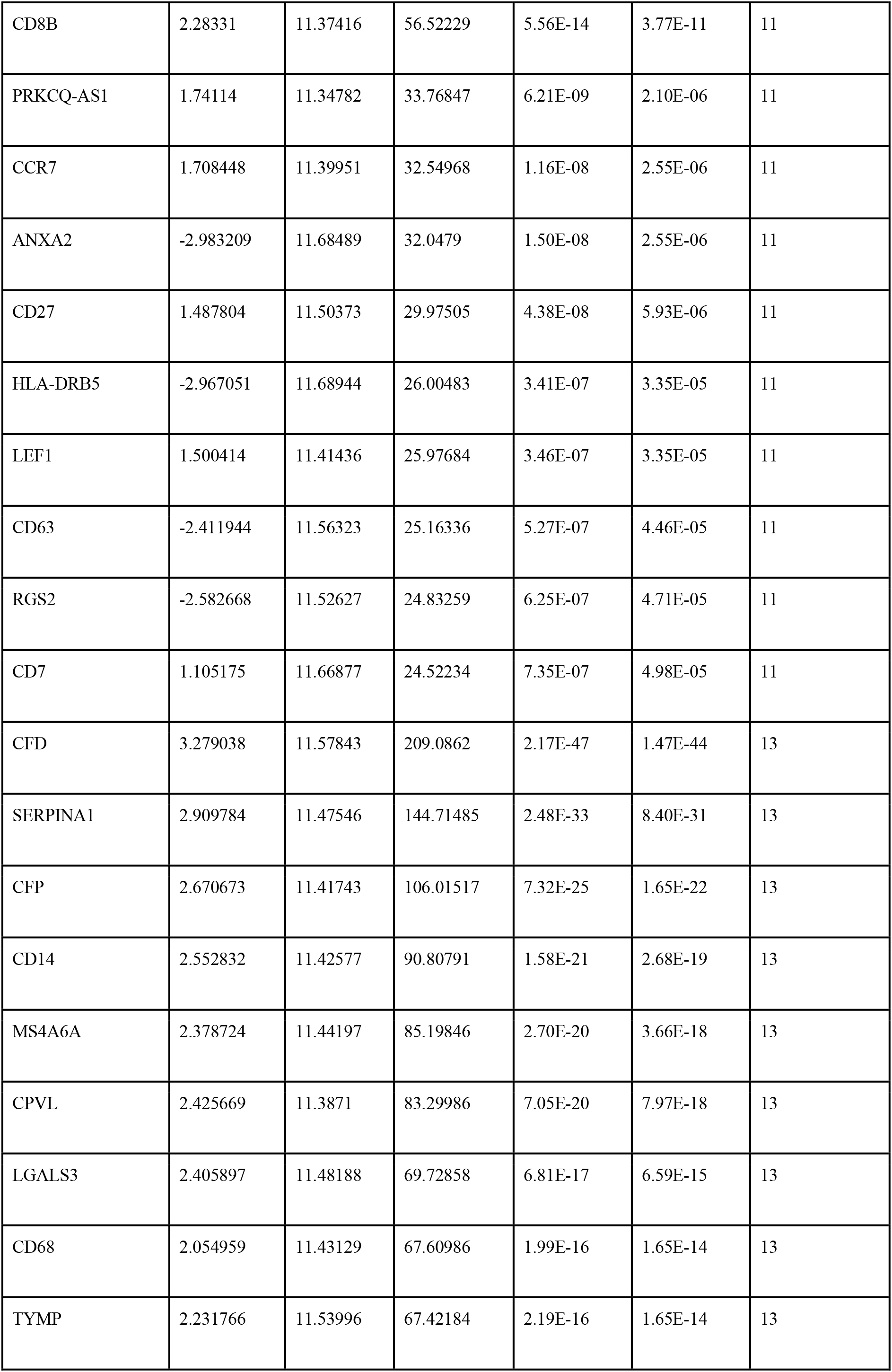
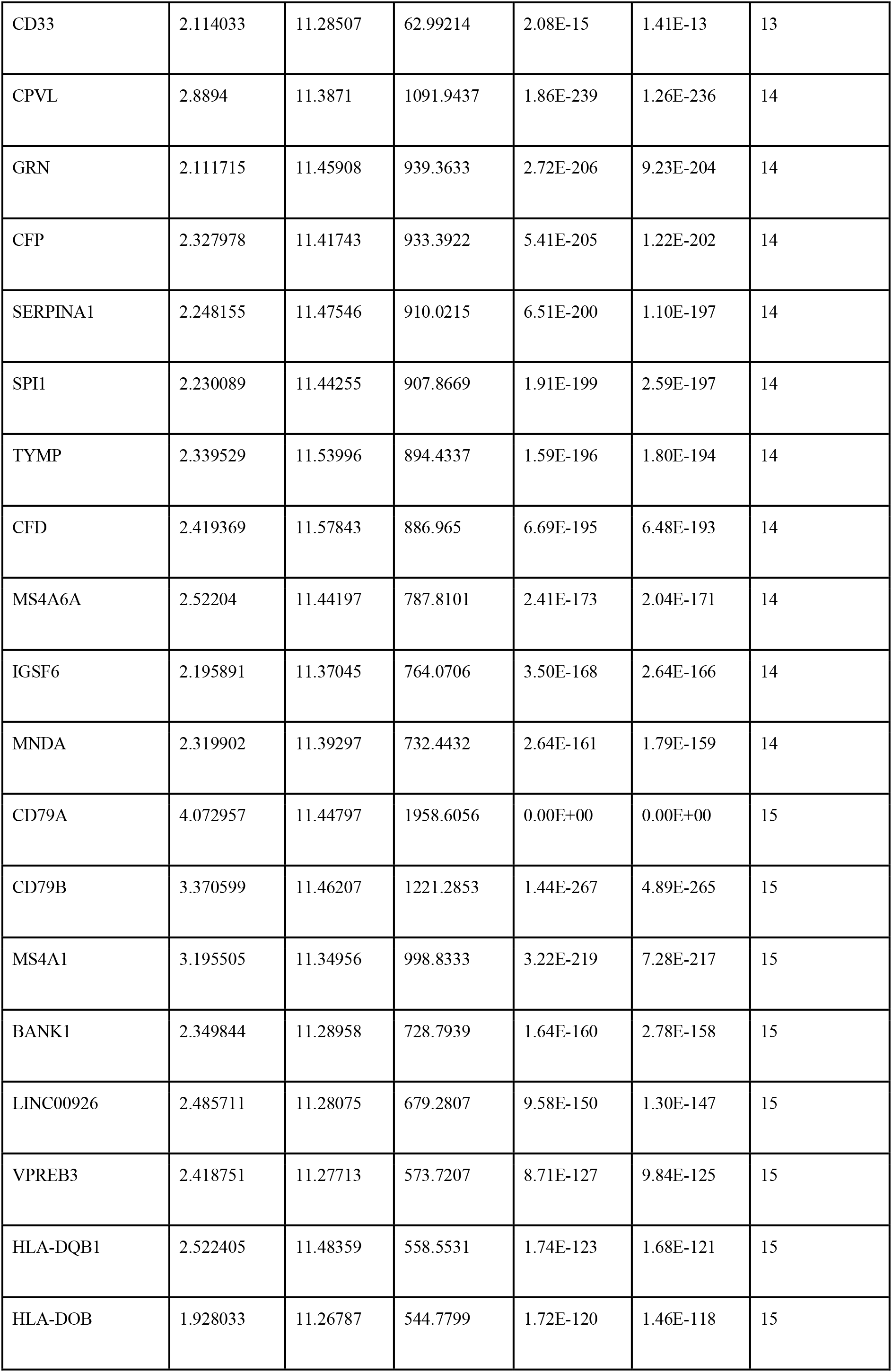
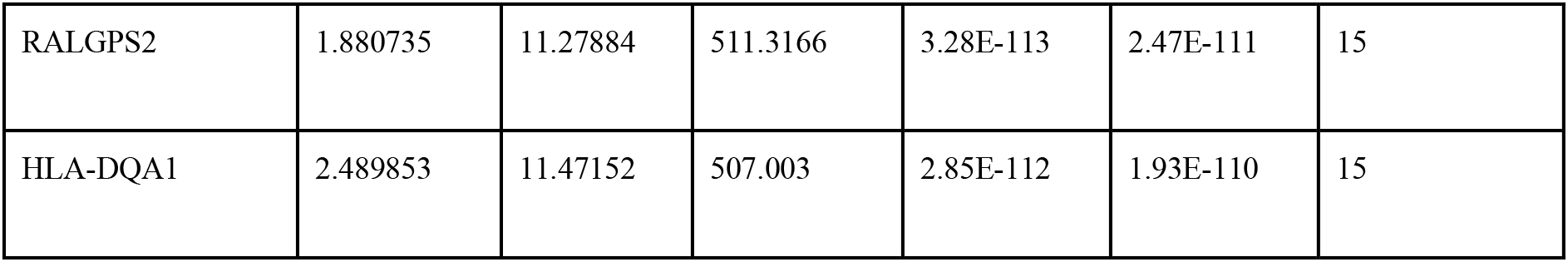
Top 10 significant genes from each cluster.

### 3.5. Cell type classification

We obtained the top significant genes for each cluster. These were positively or negatively expressed in the clusters, but only positive and significantly expressed genes were considered for classifying the cell types. For reference, we used databases such as the CellMarker database (Hu et al., 2022), The human protein atlas (*The Human Protein Atlas*, n.d., Karlsson *et al*., 2021, Uhlen *et al*., 2019), and EBI-Single cell expression atlas (*Home < Single Cell Expression Atlas < EMBL-EBI*, n.d.). The top genes were searched for cell-type-specific expression in these databases. The next table (Table 3.)shows the canonical markers chosen from each cluster to show that they are a certain type of immune cell.

**Figure 7.**
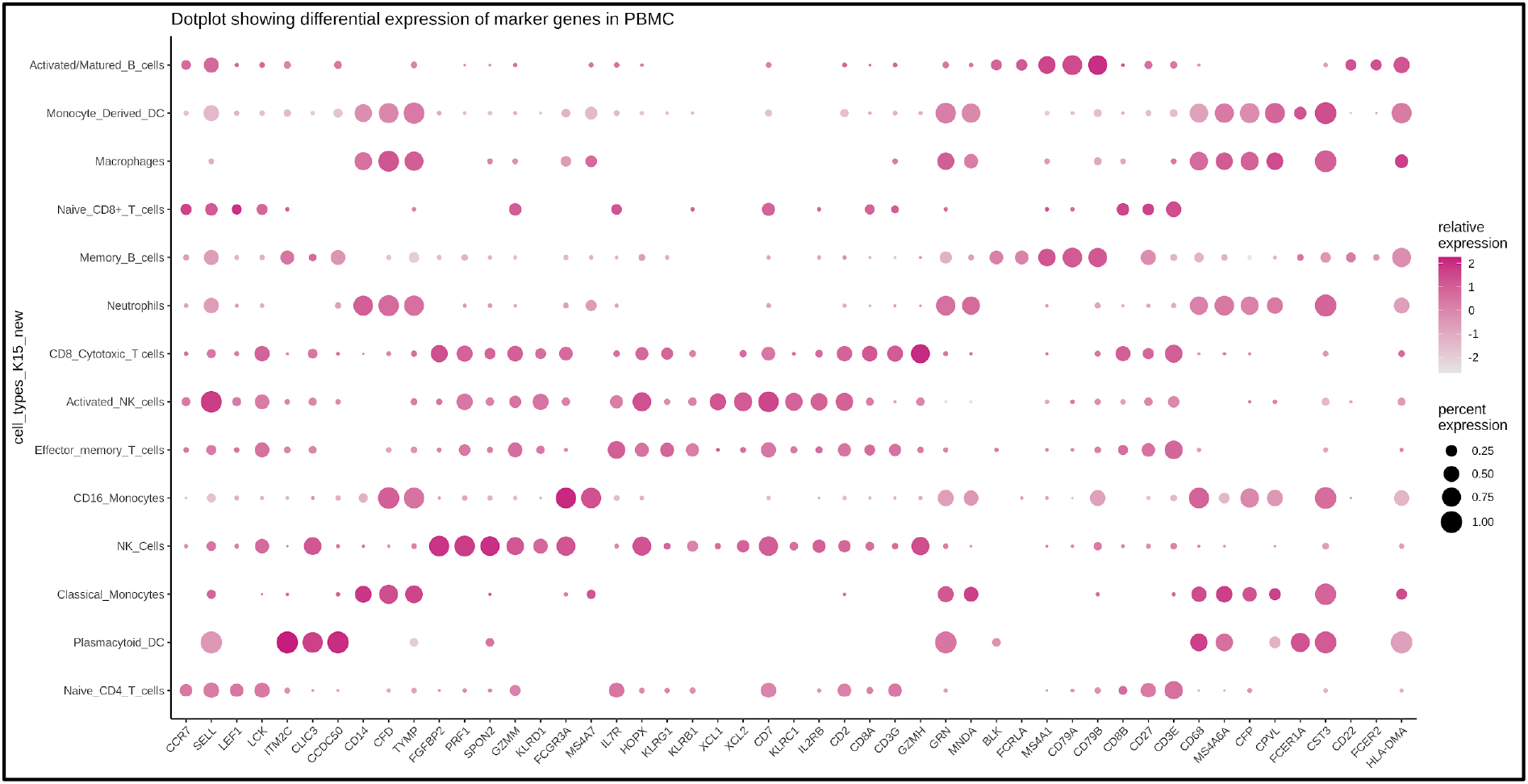
Dotplot of canonical markers and their differential expression across the clusters.

**Table 3.** Canonical markers used to identify cell types from the clusters.

| **Cluster. No.** | **Marker genes** | **Cell types assigned** |
| --- | --- | --- |
| 1 | CCR7, SELL, LEF1, LCK | Naive CD4 T cells |
| 2 | ITM2C, CLIC3, CCDC50 | Plasmacytoid DC |
| 3 | CD14, CFD, TYMP | Classical Monocytes |
| 4 | FGFBP2, PRF1, SPO2, GZMM, KLRD1 | NK cells |
| 5 | FCGR3A, MS4A7 | CD16 Monocytes |
| 6 | IL7R, GZMM, HOPX, KLRG1, KLRb1 | Effector memory T cells |
| 7 | XCL1, XCL2, CD7, HOPX, KLRC1, IL2RB | Activated NK cells |
| 8 | CD2, CD8A, CD3G, GZMH, GZMM | CD8 Cytotoxic T cells |
| 9 | CFD, GRN, TYMP, MND A | Neutrophils |
| 10 | BLK, FCRLA, MS4A1, CD79A, CD79B | Memory B cells |
| 11 | CD8B, CCR7, CD27, CD3E | Naive CD8 T cells |
| 13 | CD68, CD14, MS4A6A, CFP, CPVL | Macrophages |
| 14 | MS4A6A, HLA-DMA, FCER1A, CST3 | Monocyte Derived DC |
| 15 | CD22, MS4A1, CD79A, CD79B, FCER2 | Activated /Matured B cells |

#### Cluster 1-Naive CD4 T cells

This cluster showed a classic marker for T cells which is CD3 it also had markers for naive cells such as CCR7 and CD4 T cell marker IL7R along with other T cell markers such as SELL, LEF1, and LCK. Also, no cytotoxic T-cell markers were found; thus, this group was classified as naive CD4 T-cells.

#### Cluster 2: Plasmacytoid DC

This group had the most important marker for dendritic cells, FCER1A, along with plasmacytoid DC specific markers such as ITM2C, CLIC3, and CCDC50. These were exclusively markers for plasmacytoid dendritic cells: thus, this cluster was classified as Plasmacytoid DC. This cluster was also found closer to the B cell clusters in the low-dimensional visualizations (PCA, TSNE, and UMAP), the reason being that plasmacytoid DCs are believed to be arising from the lymphoid origin, and the same progenitors that produce B cells also produce the plasmacytoid DCs.

#### Cluster 3: Classical Monocytes

This cluster had a high expression of the typical monocyte marker CD14 and other myeloid markers such as CFD, TYMP, and MS4A6A. Thus, this cluster was classified as Classical monocytes.

#### Cluster 4: Natural Killer Cells

NK cells have the same functions as cytotoxic T cells and share the same genes responsible for killer activities in these cell types. However, the NK cells have negligible expression or mostly no expression at all for the typical T cell marker, which is CD3. Thus, NK cells can be separated from cytotoxic T cells. CD16, or FCGR3A, is also highly expressed in NK cells. This cluster lacked CD3 expression but had high levels of FCGR3A and killer genes like FGFBP2, PRF1, KLRD1, SPON2, and HOPX, as well as granzymes like GZMH and GZMM. Therefore, this group was classified as NK cells.

#### Cluster 5: CD16 Monocytes

CD16 or FCGR3A is the typical canonical marker for CD16 monocytes and this marker was expressed highly in this cluster along with other myeloid markers such as MS4A7, CD68, TYMP, and CFD. Therefore, this cluster was classified as CD16 monocytes.

#### Cluster 6: Effector memory T lymphocytes

Effector T lymphocytes are basically subtypes of memory T cells that turn into cytotoxic cells when they are activated by the right antigens. By using this method, they give the immune system “memory” against pathogens that have already been encountered. These lack the expression of the naive marker CCR7 and have a high expression of the memory marker IL7R, which was found true in this cluster. It also had other effector/killer T cell markers such as KLRG1, GZMM, HOPX, KLRB1, and CD3G. Therefore, this group was classified as effector memory T cells.

#### Cluster 7: Activated NK cells

This group showed the very first NK cell-specific marker, CD7, along with chemokines XCL1 and XCL2 and interleukin IL2RB, which are expressed in activated NK cells. The other NK cell markers were HOPX, KLRC1, SELL, KLRD1, and GZMK. Therefore, this cluster was classified as activated NK cells.

#### Cluster 8: CD8 Cytotoxic T cells

This cluster had CD8A, CD3G, and CD247 (TCR zeta) as the top significant genes, along with other killer cell markers such as GZMB, HOPX, GZMH, PRF1, and CCL4. Therefore, this cluster was classified as CD8-cytotoxic T cells.

#### Cluster 9: Neutrophils

This cluster had granulocyte-specific markers such as CFD, GRN, TYMP, and MNDA. Therefore, this cluster was classified as Neutrophils.

#### Cluster 10: Memory B cells

This cluster had B-cell-specific markers like CD79A, CD79B, MS4A1 (CD20), BANK1, antigen-presenting markers like HLA-DQA1, HLA-DQA2, and HLA-DQB1, and memory markers like PDLIM1 and BLK. Therefore, this group was classified as Memory B cells.

#### Cluster 11: Naive CD8 T cells

This cluster had the CD8 T cell marker, which is CD8B and GZMM, along with the naive markers CCR7, LEF1, and FHIT. Therefore, this group was classified as the naive CD8 T cell group.

#### Cluster 13: Macrophages

Through the differentiation of monocytes, macrophages are created. The monocytes leave the bloodstream, penetrate the infected tissue or organ, and go through a sequence of modifications to become macrophages when there is tissue damage or infection. Because monocytes are the precursors of macrophages, some markers, such as CD14 and CFD, are shared by these two cell types. It had macrophage-specific markers such as CD68, CD33, IGSF6, LY86, and FCGRT. Therefore, this group was classified as macrophages.

#### Cluster 14: Monocyte-Derived Dendritic Cells

These cells provide surface antigens for inspection by other immune system cells and are derived from monocytes under special conditions such as inflammation. These are similar to monocytes and share the same set of markers, but have DC-specific markers as well. This group had CD14 as well since these DCs originate from monocytes, but the other DC-specific markers found were CPVL, GRN, FCER1A, and antigen-presenting markers such as HLA-DMA and HLA-DQB1. Therefore, this group was classified as Monocyte-Derived DCs.

#### Cluster 15: Activated/Matured B cells

This cluster had high expression of B cell-specific markers like CD79A, CD79B, MS4A1, BANK1, maturation marker CD27, activation markers like CD22, FCER2, FCRLA, RALGPS2, and antigen-presenting markers like HLA-DQB1, HLA-DQA1, HLA-DOB, and HLA-DQA2. Therefore, this group was classified as Activated/Matured B cells.

The figure 8. shows final annotation of the clusters identified, into the cell types classified using the canonical markers. The groups and their subgroups fall closer in the plots and overlap. For example, both NK cell clusters are shown close together in the plot. The B cell subsets which are the memory B cells and activated B cells form another close group. We also observed the myeloids tightly bound together and similarly, all the T cells fall together closely in the above plots. The plasmacytoid DC although a myeloid cell is still similar to Plasma cells which are matured B cells, therefore this cell type cluster falls closer to the B cell group.

**Figure 8.**
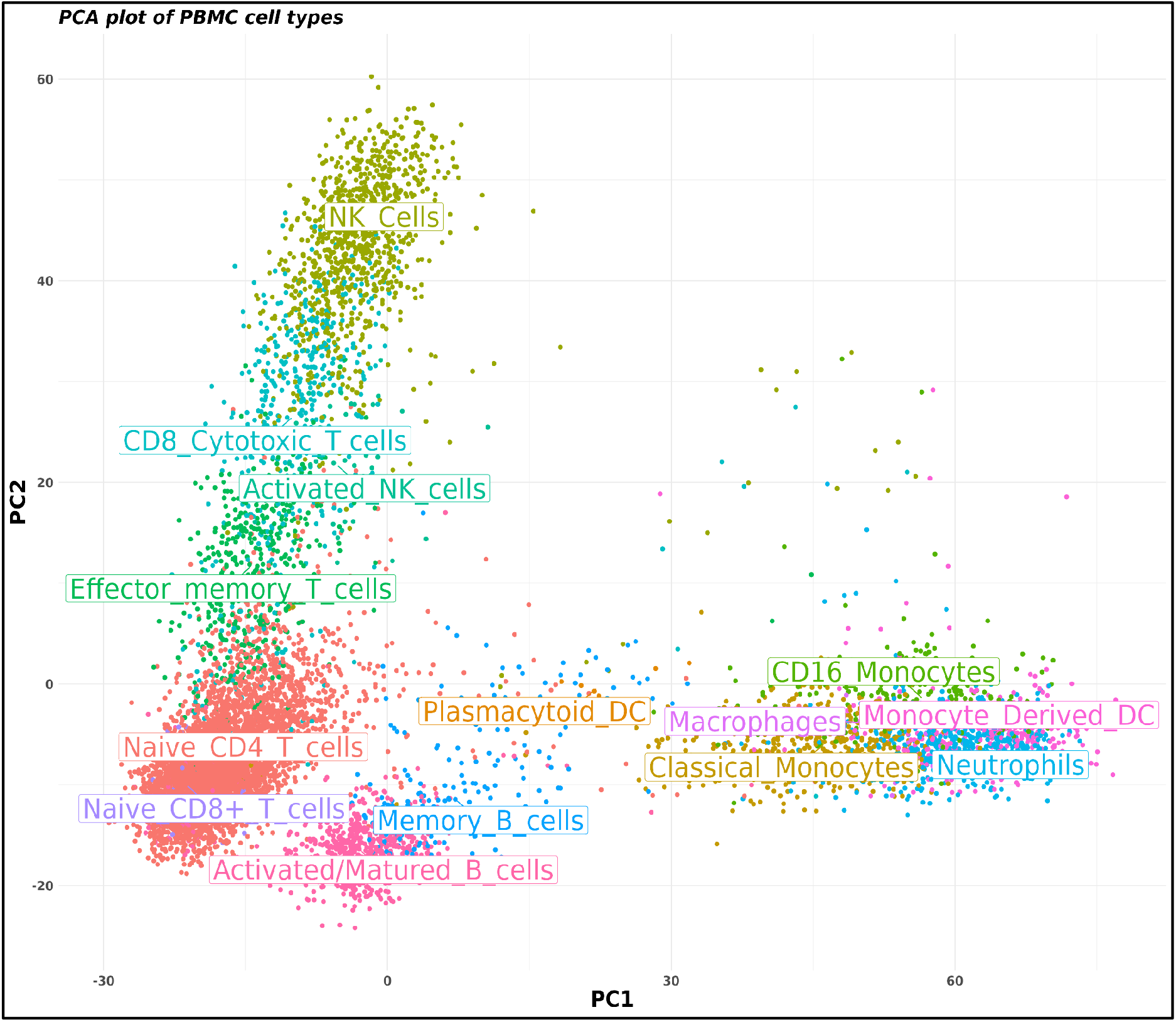
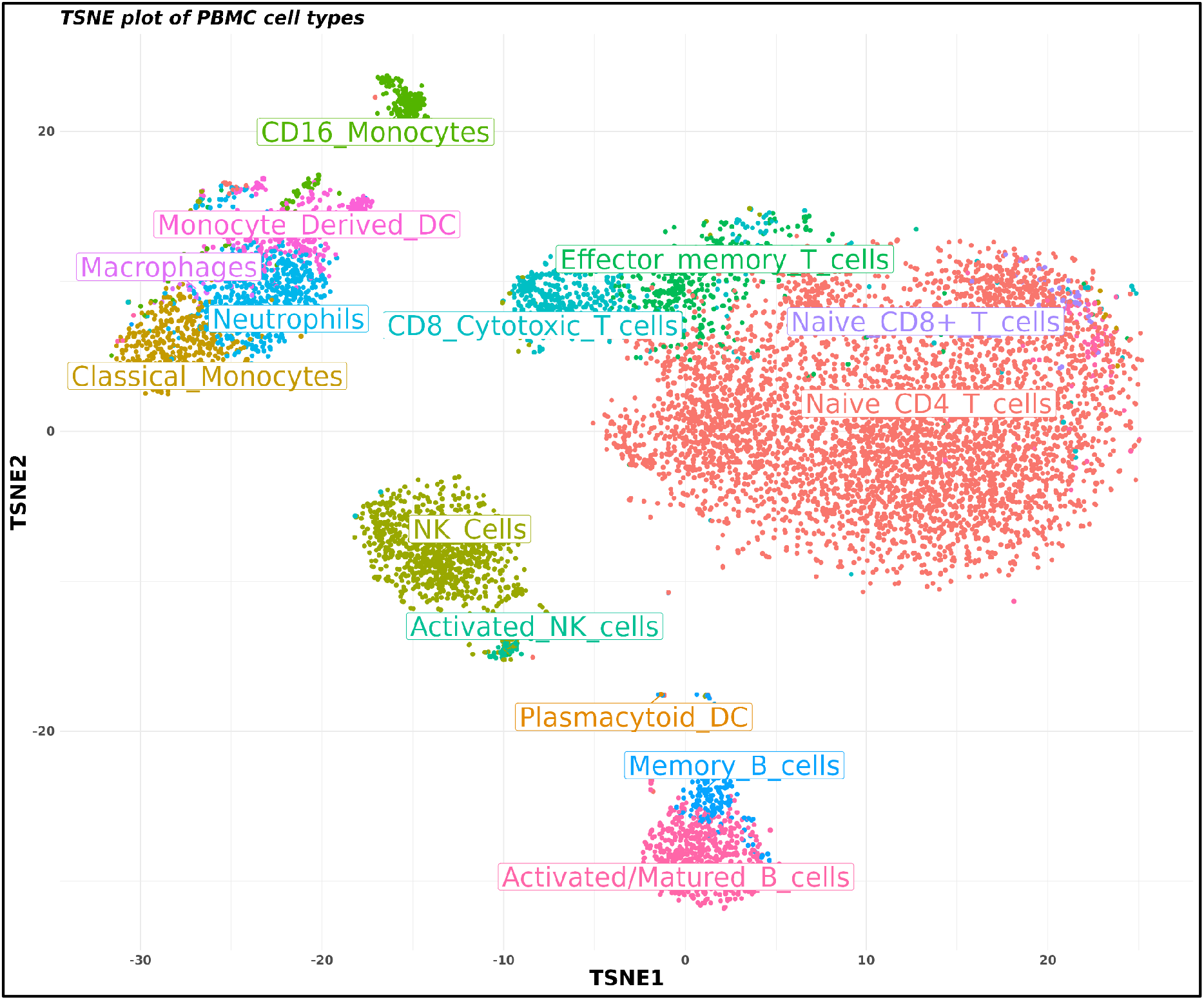
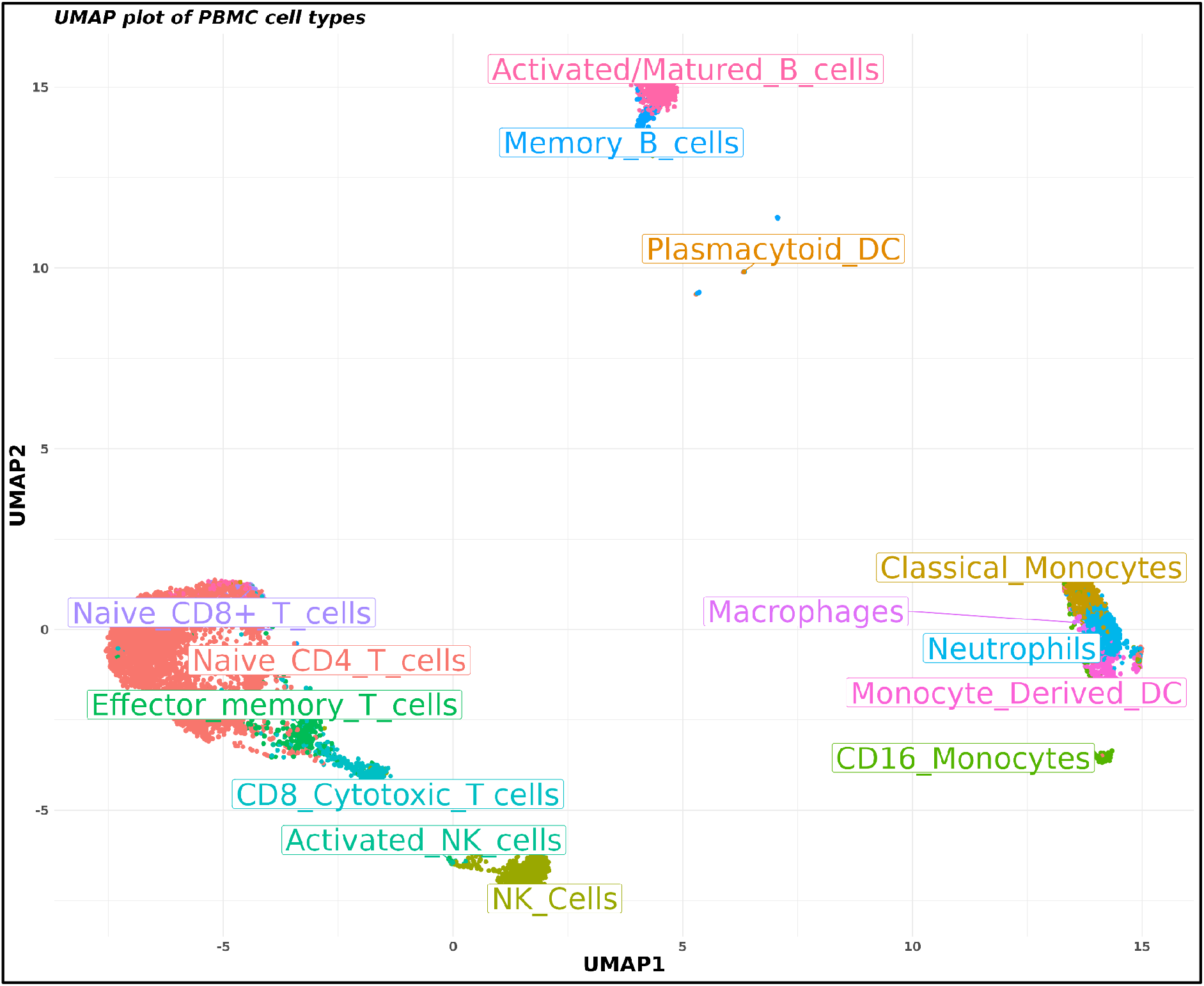
t-SNE, PCA and UMAP plots showing the cell types identified in the PBMC data. Note that the T cells are grouped together; similarly, the B cells are also grouped together; the CD16 monocyte is a distinct cluster, as is the one with NK cells. All the myeloid cells, such as DCs, monocytes, macrophages, and neutrophils, are grouped closely. Another thing that stands out is that plasmacytoid DCs are close to B cells, which makes sense because plasmacytoid DCs are similar to plasma cells.

### 3.6. Construction of the Signature Matrix

The signature matrix was built after classifying the clusters as the types of cells with genes that were highly variable across the cells. We had 14 cell types and 678 genes and constructed the signature matrix by the method mentioned above. The reference file was prepared to have each cell labelled as the cell type it belonged to. Non-hematopoietic genes were set to be filtered from the resultant matrix. We obtained a signature matrix with 14 columns representing 14 cell types and 275 genes. The matrix had values for canonical markers the highest for their respective cell types (up to thousands) and 1 everywhere else. For example, The B cell marker, MS4A1, was the highest in the two B cell subtypes (memory B cells and activated/matured B cells) but the least (1) for other cell types. Similarly, the memory markers for B cells were high compared to those in activated B cells, but they were the least, i.e., 1 for every other cell type. The same was true for other cell types and subtypes as well.

Figure 9. shows a heatmap for the differential expression of the canonical markers in the custom signature matrix. It also shows the hierarchical clustering of the cell types. Heatmap shows the myeloid group separated from the lymphoid group. Under the Myeloid group fall the monocytes, DCs and macrophages. It also shows the macrophages and Monocyte DCs grouped together which is correct as they both are antigen presenting cell and originate from monocytes. Similarly, the neutrophils and monocytes grouped together since they both are phagocytes that act against infections. They also share some of the markers due to similar type of functions, thus, this grouping holds. The plasmacytoid DCs in the myeloid group but separate from other myeloid cell types. This is because although myeloid, the plasmacytoid DCs have similarities with the plasma cells (or matured B cells), they fall in the myeloid group because they have antigen-presenting markers, a characteristic of dendritic cells and other myeloid and dendritic cell markers.

**Figure 9.**
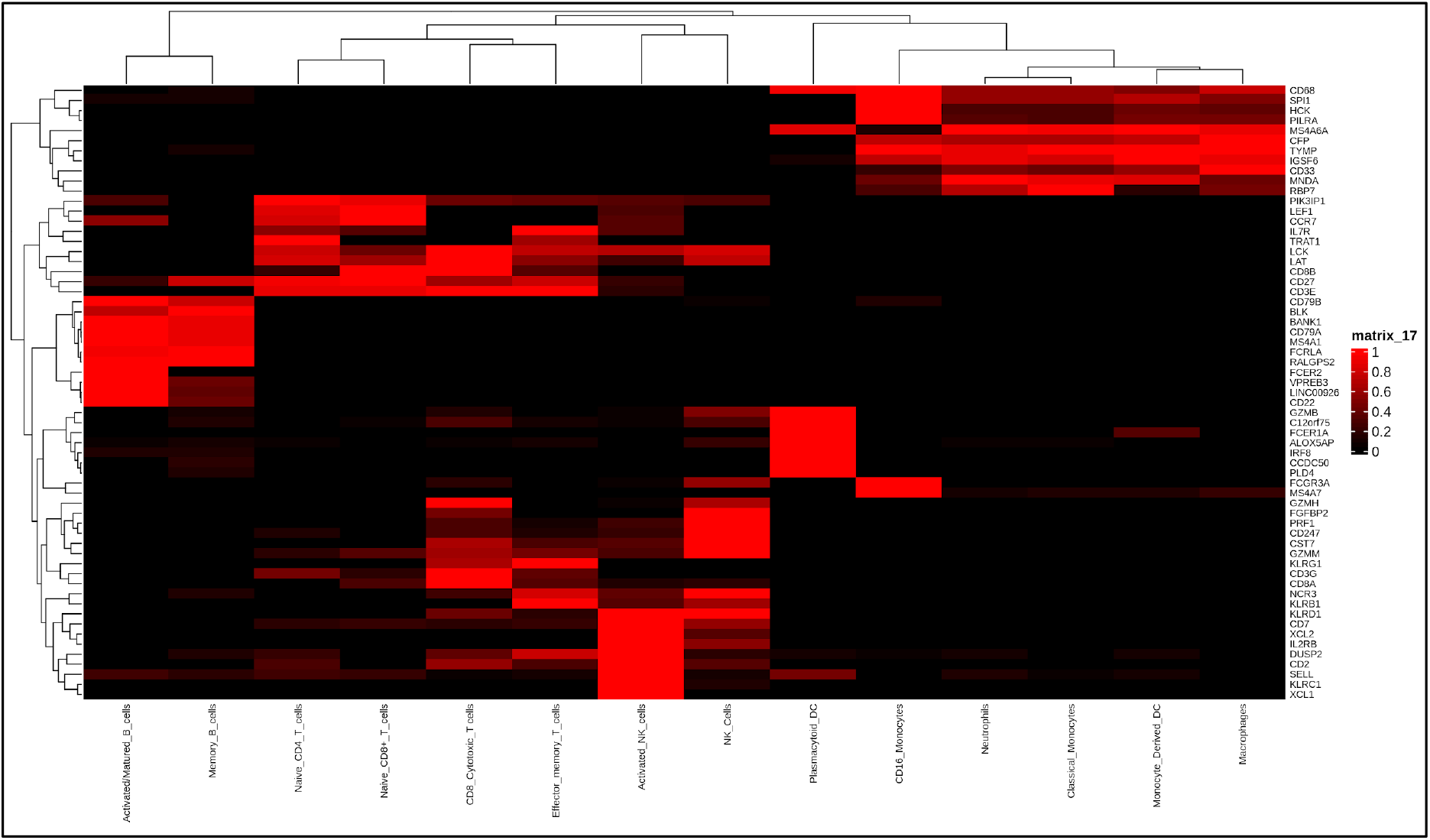
Heatmap showing the differential expression of the canonical markers in the custom signature matrix.

In the lymphoid group, we see three different sections. One is for B cells described earlier, the otehr is T cells and one more which is the NK cells group. The NK cell group has both the NK cell subtypes which holds true for obvious reasons. The T cell group has the cytotoxic and effector T cells grouped together and both the naive groups together. This grouping is true because the effector T cells are also capable of killer functions just like the cytotoxic T cells, thus they have similar expression profiles and many common markers. The naive group has both the naive cell forms of T cells.

SOUP selected 678 genes when the gene selection was performed using default parameters. However, the number of selected genes may be increased or decreased when the parameters such as “threshold” and “sumabs” are changed in the gene selection step. The “threshold” is the cutoff for the Gini index of DESCEND. Fewer genes are picked when the threshold is higher. Similarly, “sumabs” is a measurement of the sparsity of genes in SPCA, between 1/sqrt(number of genes) and 1. Smaller values result in sparser results, hence fewer selected genes.

The 678 genes are able to segregate the data into 8 distinct clusters. If, by changing the parameters in the gene selection step, the number of highly variable genes is increased or decreased, the number of groupings may change when identified by an elbow plot. However, it won’t really change the overall grouping structure in the 3 visualizations (PCA,t-SNE, UMAP). Thus, to sum up, the number of genes selected for further clustering the cells, play a crucial role in cluster segregation.

Since there were more than 8 groupings in the visualization plots (PCA, t-SNE, UMAP). These groupings could be cell subtypes, which is why the clustering was further continued till K15, after that, there were clusters with less than five cells. Therefore, it was stopped at k15.

## 4. Conclusion

The single-cell data helps to identify the various cell types and subtypes accurately, with a precise set of markers and genes for each cell type, since it has a relatively higher resolution than bulk RNA-seq data.

The CIBERSORTx toolkit uses large-scale tissue transcriptome profiles to look at the abundance of different cell types and how genes are expressed differentially in them. With the “custom signature matrix construction” utility, the genetic markers of many cell subsets can be learned out of a smaller number of biological specimens or samples using single-cell or bulk-sorted RNA sequencing data. Then, these signatures are used to untangle the bulk tissue transcriptomes to find out the abundance of each type of cell.

With the help of the scRNA-seq PBMC dataset, we made our signature matrix after finding out about 14 different types and subtypes of immune cells. After hierarchical clustering, each cell type was assigned in the right group because it had high expression of its canonical markers. This signature matrix needs to be validated on positive and negative controls (bulk RNA-seq), and then it can deconvolve the traditional RNA-seq cancer datasets.

## Appendix

### Supplementary files

- PBMC clusters obtained after differential expression analysis showing 678 genes including differential and significant expression of marker genes, in different clusters.

- Cluster_1
- Cluster_2
- Cluster_3
- Cluster_4
- Cluster_5
- Cluster_6
- Cluster_7
- Cluster_8
- Cluster_9
- Cluster_10
- Cluster_11
- Cluster_13
- Cluster_14
- Cluster_15
- Custom_Signature_matrix : Custom Signature matrix of 14 PBMC immune cells created using CIBERSORTx

## Abbreviations

Cox: Reid profile-adjusted likelihood (CR)
CV: cross validation
DCs: Dendritic cells
DESCEND: deconvolution of single-cell expression distribution
EBI: European Bioinformatics Institute
FM: Ferragina-Manzini
GEMs: Gel beads in emulsion
GLM: Generalized linear model
HVG: Highly Variable genes
LM22: Leucocyte Matrix 22
NGS: Next Generations Sequencing
NK cells: Natural Killer cells
PBMC: Peripheral Blood Mononuclear Cells
PCA: Principal component analysis
QL: Quasi likelihood
scRNA-seq: Single cell RNA sequencing
SE: squared error
SOUP: semisoft clustering with pure cells
SPCA: Sparse Principal component analysis
SVM: Support vector machine
SVR: Support vector regression
T cells: T lymphocytes
T-regs: Regulatory T Cells
t-SNE: T-distributed stochastic neighborhood embedding
TMM: trimmed mean of M values
TPM: Transcripts Per Kilobase Million
UMAP: uniform manifold approximation and projection
UMI: Unique Molecular Identifier

